# Critical Assessment of a Structure-Based Pipeline for Targeting the Long Non-Coding RNA MALAT1

**DOI:** 10.1101/2025.11.13.688270

**Authors:** Riccardo Aguti, Mattia Bernetti, Andrea Cavalli, Matteo Masetti

**Author notes:** **Corresponding Author:** Matteo Masetti, Mattia Bernetti.

## Abstract

Long non-coding RNAs (lncRNAs) are increasingly recognized as druggable targets due to their conserved secondary/tertiary structures and regulatory roles in disease. A prototypical example is the MALAT1 triple helix, whose stability supports transcript persistence and is implicated in oncogenesis. Here, we evaluate the ability of a structure-based drug discovery (SBDD) pipeline, integrating molecular dynamics (MD), pocket analysis, ensemble docking, and diverse scoring functions, to capture the binding behavior of 21 congeneric diminazene-based ligands targeting MALAT1. Conformational ensembles were generated using both conventional MD and Hamiltonian Replica Exchange MD, revealing two potential binding sites. Ensemble docking with AutoDock GPU and rDock across representative RNA conformations, followed by rescoring with force-field and machine-learning-based scoring functions, led to the identification of a binding mode with the best agreement across the series. Principal component analysis of interaction fingerprints within clustered poses was used to explain the experimentally observed affinity trends. Our findings highlight the promise and limitations of current SBDD pipelines for flexible RNA targets and offer a benchmark for future improvement in RNA-focused drug discovery.

## Introduction

The organization of the eukaryotic genome is highly complex, with nearly 98% of the human genome consisting of non-coding sequences.(*1, 2*) Once regarded as “junk DNA”, these non-coding DNA regions are now recognized as a rich repository of functional elements, including various classes of non-coding RNAs (ncRNAs).(*3–6*) The extent of functionality within these non-coding sequences remains a subject of ongoing debate.(*7–9*) Nevertheless, findings from the Encyclopedia of DNA Elements (ENCODE) project show that approximately 80.4% of the full genome is involved in biochemical processes such as chromatin organization, histone modification, and transcriptional regulation.(*10*)

Non-coding RNAs are further classified based on their size. Short ncRNAs, with less than 200 bases in length, include tRNAs, rRNAs, miRNAs, and snoRNAs.(*11*) Conversely, long non-coding RNAs (lncRNAs), exceeding 200 nucleotides in length, constitute a diverse class of regulatory molecules.(*12*) lncRNAs often exhibit greater conservation in their secondary and tertiary structures compared to their primary sequences.(*13*) This structural conservation underscores their functional importance, yet their flexibility complicates experimental structure determination, even with emerging techniques such as Cryo-EM.(*14, 15*) Being the most prevalent class of ncRNAs, lncRNAs are involved in critical biological processes such as epigenetic regulation, transcriptional control,(*16*) X chromosome inactivation,(*17, 18*) genomic imprinting,(*19, 20*) and cell development.(*21*) Consequently, their dysregulation is linked to various diseases, including cancer.(*22*)

A key example of the pharmacological relevance of lncRNAs is the metastasis-associated lung adenocarcinoma transcript 1 (MALAT1).(*23*) Its overexpression has been related to various cancers,(*24*) and its knockdown has been shown to reduce oncogenic processes,(*25, 26*) among other phenotypic effects.(*27–29*) However, the precise mechanisms and interaction partners underlying MALAT1’s regulatory functions remain an active area of research.(*30*) MALAT1 is a promising target for small-molecule therapeutics due to the presence of a well-characterized 3’-terminal triple helix, which plays a critical role in transcript stability (**Fig. 1A**). This structure, functionally characterized in cell-based assays(*31*) and resolved through X-ray crystallography,(*32*) protects MALAT1 from degradation by sequestering its adenine-rich 3’-tail through base pairing with a uridine-rich stem loop. Beyond its contribution to overall stability, recent research suggests that local conformational dynamics can affect the transcript protection, as increased structural flexibility or unfavorable ionic concentration hinders its integrity and leads the RNA to be vulnerable to exonucleolytic degradation.(*33*) Notably, small molecules can influence both the global stability and the conformational landscapes of RNA,(*34–37*) further highlighting the potential for targeted therapeutic intervention.

**Figure 1.**
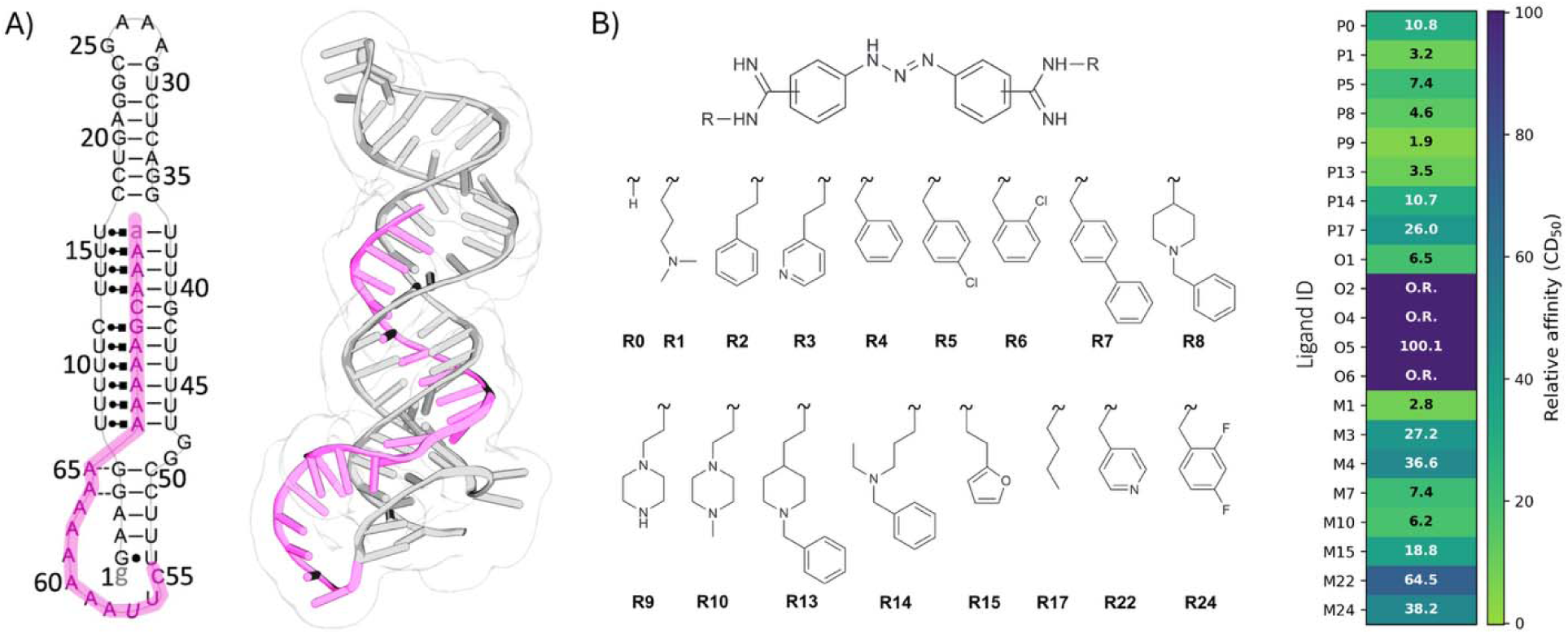
The MALAT1 RNA triple helix and the diminazene series of ligands. A) Secondary (left) and tertiary (right, PDB ID: 4PLX) structure of the MALAT1 triplex, with the poly(A) tail highlighted in light purple. B) The dataset of small molecule ligands with diminazene scaffold used in this work, and corresponding experimental relative binding affinities in µM (right panel). Relative affinity is expressed as CD_50_ (small molecule concentration needed to achieve 50% competitive displacement); molecules labeled as out of range (O.R.) indicate small molecules with CD_50_ values outside of the range that lead to minimal dye displacement in the assays (>500 µM). The position of the substituents on the central scaffold (ortho, O, meta, M, or para, P), and the ID of the substituent compose the ligand ID.

Structure-based drug discovery (SBDD) is routinely exploited in both academia and industry to accelerate the identification and optimization of bioactive compounds.(*38*) However, most computational SBDD tools have been developed for proteins, as the relevance of RNA as a pharmaceutical target has only recently gained widespread recognition.(*39–41*) Consequently, their application in the context of RNA targeting has received limited exploration.(*40–50*) Notably, the performance of conventional docking tools and their counterparts specifically developed for RNA remains uncertain. For example, while the RNA-specific docking program rDock outperformed AutoDock4 in pose prediction,(*48*) broader benchmark studies have generally found protein-specific tools, including AutoDock GPU and AutoDock VINA among others, to perform better overall.(*49*) Furthermore, a more recent evaluation of the enrichment capabilities of different docking programs in virtual screening campaigns revealed that none of the tested methods performed consistently well across all relevant metrics.(*50*) Importantly, these benchmarking studies have not investigated the role of computational methods in generating binding-competent conformations of the RNA targets or identifying suitable pockets for small-molecule recognition. The disclosure of a library of 21 congeneric diminazene-based ligands targeting the MALAT1 triple helix, along with their experimentally determined affinities (**Fig. 1B**),(*51*) provides an ideal test case for assessing the current capabilities of computational SBDD approaches to RNA targets. In particular, MALAT1 exemplifies the complexity of a real-world scenario, where the conformational landscape is highly dynamic and neither the binding mode nor the exact binding site is known a priori.

In this work, we evaluate the applicability of SBDD frameworks based on molecular docking and Molecular Dynamics (MD) simulations to the MALAT1 triplex (**Fig. 2**). Specifically, we use conventional MD(*52*) and Hamiltonian Replica Exchange (HREX)-MD(*53*) to thoroughly explore the RNA’s conformational space(*54*) and identify potential small-molecule binding sites. Then, we perform ensemble docking of the 21-ligand library against representative MALAT1 conformations using AutoDock GPU(*55*) and rDock.(*56*) Finally, we assess the resulting poses with a range of force field- (AutoDock,(*57*) AutoDock Vina,(*58*) rDock(*56*)) and machine learning-based (AnnapuRNA,(*59*) SPRank(*60*)) scoring functions to determine the ability of each protocol to recapitulate the experimentally observed variation in binding affinity and binding mode consistency across the library. Our findings identified two persistent and ligand-accessible binding sites, with one site featuring a cluster of poses with a distinctive and consistent binding mode of the central scaffold, providing insight into ligand affinity trends. While the overall accuracy of the considered scoring functions remained limited, our integrated approach uncovered meaningful structure–affinity trends and revealed key challenges and opportunities in RNA-targeted drug discovery.

**Figure 2.**
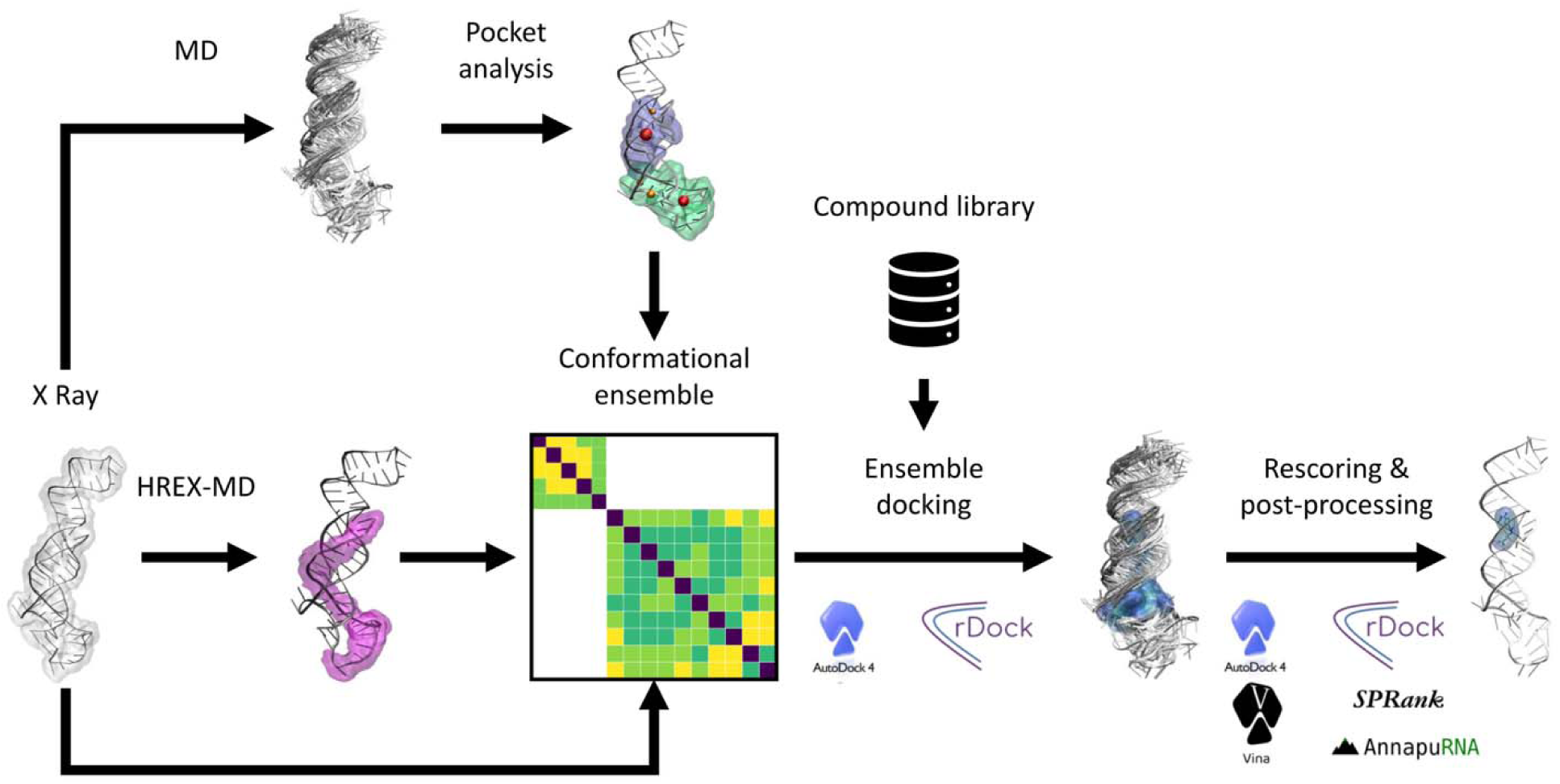
The computational pipeline established in this work. Starting from the crystal structure, standard and advanced MD simulations are employed to identify promising binding regions and to explore the RNA conformational dynamics, respectively. The information is integrated to generate a conformational ensemble of the identified binding regions, which is then used for ensemble docking of the diminazene library of compounds; ligand poses at this stage are produced using the AutoDock 4 and rDock software. Finally, the binding poses are rescored using diverse scoring functions (AutoDock 4, rDock, Vina, SPRank, and AnnapuRNA), and post-processing of the outcomes allows rationalization of the results.

## Methods

### System Preparation

To perform molecular dynamics (MD) simulations, the RNA target structure (PDB ID: 4PLX)(*32*) was modeled using the Amber force field for nucleic acids, comprising the Amber f99SB force field(*61*) and the bsc0(*62*) and XOL3 refinements(*63*).

The system was solvated using the four-point OPC water model,(*64*) while potassium and chloride ions were added to neutralize the system and achieve a physiological salt concentration of 0.15 M using the Joung and Cheatham ion model.(*65*) Subsequently, the system underwent energy minimization followed by equilibration in both the NVT and NPT ensembles, with a total simulation time of 1.2 ns (600 ps for each ensemble) with a timestep of 2 fs. Covalent bonds involving hydrogen atoms were constrained using the LINCS algorithm,(*66, 67*) while long-range electrostatic interactions were treated with the Particle Mesh Ewald (PME) method,(*68, 69*) using a cutoff distance of 1.2 nm.

For conventional MD simulations in the NPT ensemble, the final equilibrated system was used as the starting point. For Hamiltonian Replica Exchange (HREX) simulations,(*53*) the trajectory frame with volume closest to the mean from the second half of the NPT equilibration was used as initial configuration for production in the NVT ensemble. This choice mitigated artifacts from box size fluctuations that could lead to abnormal pressure values during subsequent HREX MD simulations performed under NVT conditions.

### Molecular Dynamics Simulations

Following system preparation and equilibration, production MD simulations were conducted on the MALAT1 RNA system, namely conventional MD and HREX MD.(*53*) Conventional MD simulations were run in the NPT ensemble at 300 K and 1 bar, maintained by the V-rescale thermostat(*70*) and C-rescale barostat,(*71*) respectively. Three independent replicates, each lasting 500 ns long, were performed.

To enhance the conformational sampling of the RNA molecule, we employed the HREX scheme, which involves simulating multiple replicas of the systems with modified Hamiltonians where charges, Lennard-Jones parameters, and dihedral potentials of a predefined region of the system are gradually scaled.(*53*) In this study, 16 replicas were used with the scaling parameter A ranging from 1 to 0.7, applied specifically to residues 55 to 76 of the poly(A) in the triple helix (**Fig. 1A**). HREX simulations were run in the NVT ensemble at 300 K using the V-rescale thermostat.(*70*) Each replica was simulated for approximately 96 ns, with exchange attempts every 240 steps. The trajectory from the “cold” replica (A = 1) was extracted and processed for further analysis.

The conformational variability of the poly(A) tail of the triple helix was inspected in both the conventional and HREX MD simulations, focusing on the “coldest” replica in the latter case, by computing the RMSD, aligning and computing on the RNA heavy atoms, and eRMSD, based on nucleotide g-vectors, with respect to the starting structure.(*72*) While the RMSD measures the overall structural deviation, the eRMSD is specifically designed for RNA and reflects base-pairing rearrangements, with values above 0.7–0.8 typically indicating discrepancy in the base-pairing pattern with respect to the reference.(*72*)

### Pocket Identification

The three trajectories from conventional MD of the RNA were aggregated and analyzed with the Pocketron algorithm to identify dynamic pocket formation and communication.(*73, 74*) Pocketron, included in the BiKi Life Sciences software package,(*75*) tracks the dynamic evolution of binding pockets during MD simulations, using the NanoShaper(*76, 77*) algorithm to detect pockets on a frame-by-frame basis. The approach leverages solvent-excluded surface calculations with two probe sizes, and we used here the default radii of 1.4 Å and 3.5 Å for the smaller and larger probes, respectively. To focus on pockets with potential relevance for small-molecule binding and long-range allosteric communication, only cavities larger than the volume of three water molecules were tracked. This filtering step excludes highly transient or solvent-exposed pockets that are unlikely to accommodate ligands. The analysis allows identifying pockets exhibiting the largest volume, highest persistence, and significant long-range communication.

### Conformational Ensemble Preparation

The HREX-MD simulation was analyzed to extract a representative ensemble of conformations for subsequent docking studies. The conformational variability of the regions identified as promising binding sites, based on the pocket analysis on the conventional MD runs, was evaluated in the “coldest” replica (A=1) of the HREX. We first determined the RMSD, aligning and computing on the RNA heavy atoms, and eRMSD, based on nucleotide g-vectors, with respect to the starting structure.(*72*)

Principal Component Analysis (PCA) was performed via the scikit-learn python package,(*78*) using as input features the g-vectors of residues comprised in the two selected binding regions. Finally, to identify representative conformations of the two sites from the HREX-MD simulation, we applied the Quality Threshold (QT) clustering algorithm (*79*) to the eRMSD matrix of the frames from the coldest replica, using a cutoff of 0.7 to group similar structures and extracting centroids from each of the identified cluster. We then retained cluster centroids with eRMSD > 0.7 with respect to all other cluster centroids in order to maximize the structural variability of the ensembles. The procedure was repeated separately for the two binding regions identified.

### Docking Pose Generation

Docked poses of the 21 diminazene analogs were generated using two software packages, namely AutoDock GPU(*55*) and rDock, with default settings(*56*) Both software require dedicated preparation of the biomolecular target and the ligands, using a united-atom representation with explicit hydrogens on polar heavy atoms only. A notable difference between the two programs is the grid definition for docking. In both software, the grid was centered on the center of mass of the heavy atoms of the residues comprised in the binding sites. In AutoDock GPU, a rectangular grid needs to be defined (**Fig. S1**), for which we employed 110 × 110 × 130 points for the first site and 110 × 80 × 80 points for the second one, with a grid spacing of 0.3 Å between points. These dimensions were chosen to ensure complete coverage of the binding regions identified through Pocketron. Differently, rDock employs a two-sphere approach for grid generation. Here, we used a radius of 17 Å for both sites, with a small probe of 1 Å and a large probe of 17 Å. This aimed at creating a spherical grid encompassing about the dimensions of the AutoDock one, to ensure comparable volumes allowed for pose generation via the two docking software. For each of the 21 ligands, we generated 250 poses with each software.

### Pose Rescoring

To evaluate the generated ligand binding poses on the RNA target, we applied multiple scoring functions, encompassing both force field-based and machine learning-based approaches. Specifically, as force-field-based methods, we employed the scoring functions from AutoDock,(*57*) rDock,(*56*) and Vina.(*58*) On the machine learning side, we used the recently developed AnnapuRNA(*59*) and SPRank(*60*) scoring functions. AnnapuRNA leverages a coarse-grained representation of RNA and ligand pharmacophores, which is used to determine RNA-ligand interaction scores using a trained ML model. SPRank refines knowledge-based pairwise potentials iteratively against experimental RNA-ligand structures to discriminate correct poses.

The Vina, AnnapuRNA, and SPRank scoring functions can be directly applied to the output poses from AutoDock GPU and rDock after minor formatting conversion. However, artifacts arose when rescoring some of the poses generated by rDock with the AutoDock GPU scoring function, evidenced by high score values. To address this issue, the grid box for both binding regions was extended to 130 × 130 × 130 points in the three dimensions during the rescoring phase. For consistency, also the poses generated by AutoDock GPU rescored using this grid definition.

### Interaction Fingerprints

To characterize RNA-ligand interactions in the generated binding poses, we computed interaction fingerprints via the fingeRNAt(*80*) python package. The tool analyzes complexes between nucleic acids and ligands to generate binary vectors indicating presence or absence of specific non-covalent interactions, namely hydrogen bonds, halogen bonds, lipophilic interactions, cation-anion, π-stacking, π-cation, and π-anion. The software inspects the residues in nucleic acid structures using geometric criteria, and the analysis supports three modes: SIMPLE, for detecting basic atomic contacts; PBS, for distinguishing interactions involving phosphate, base, and sugar moieties of nucleic acids; FULL, for classifying the interactions into specific categories. Herein, we applied fingeRNAt to the RNA-ligand binding poses using the FULL method. For every pose, histograms of interaction occurrences were computed at distances ranging from 2 to 8 Å with 80 bins. Additionally, PCA was applied on the interaction fingerprints, using bin populations as input features.

### Clustering of the binding poses

Cluster analysis of the binding poses was achieved through the Quality Threshold (QT) clustering algorithm(*79*) applied to the minimum distance RMSD matrix(*81*) with a threshold of 5 Å. The RMSD matrix was computed focusing on the common diminazene scaffold of the 21 compounds. For proper clustering of the generated ligand binding poses, structural symmetry must be taken into account. To this end, RMSD values between pose pairs were computed on the Cartesian coordinates of the ligand scaffold both directly and after applying symmetry transformations,(*82*) and the minimum RMSD value was retained. This was particularly important for para-substituted ligands with symmetrical side chains (**Fig. S2**).

## Results and Discussion

### Sampling the Conformational Space of the MALAT1 Triple Helix

Generating an ensemble of drug-target conformations by MD simulations for subsequent virtual screening campaigns is a well-established SBDD approach.(*83*) Motivated by the need to explore MALAT1’s conformational space and identify suitable binding pockets, we carried out simulations starting from the available X-ray structure (PDB ID: 4PLX).(*32*) Specifically, we performed three independent conventional MD simulations alongside a single HREX-MD simulation. It is indeed well known that standard MD can suffer from limited sampling,(*84*) and HREX-MD was employed to enhance the coverage of accessible conformations. The triplicate conventional simulations then served as a baseline for quantifying the benefit of this enhanced sampling protocol.

**Figure 3A** illustrates the conformational dynamics of the poly(A) tail under conventional MD sampling. In particular, the RMSD analysis reveals significant structural fluctuations, especially in Run 1, where RMSD values reach up to 8 Å. In contrast, the eRMSD timeseries remain more consistent across all replicates, with values centered around 0.7, and only occasionally approaching 1 in Run 3. This suggests that the high RMSD values observed do not necessarily correspond to significant base pairing rearrangements, emphasizing the importance of eRMSD as a more tailored metric for capturing relative changes in nucleic acid structure. Furthermore, the PCA plot, color-coded by replica, shows that even relatively short simulations access a distinct conformational space in the three runs. As a comparison, **Figure 3B** demonstrates the effectiveness of HREX-MD in sampling the conformational landscape of MALAT1 more exhaustively, as evidenced by the corresponding eRMSD timeseries, reaching values up to 1.2. The improved sampling can be better appreciated through dimensionality reduction via PCA (**Fig. 3C**), which shows that the HREX-MD not only fully covers the PCA space spanned by the three conventional MD runs but also explores previously uncharted regions. By analyzing and visually comparing the centroid structures, we observe that, compared with the conventional MD, the HREX simulations promote greater mobility of the poly(A) residues, particularly in the lower region where the triple helix wraps around the duplex and inserts to form the triple pairing. In this region, where the residues are less constrained, larger displacements are detected due to lost interactions involving particularly residues U56–U57, A60–A61, and U52–U53. This enhanced flexibility also reveals a new cluster of conformations sampled only in the HREX simulations, in which the lower region folds back on itself, bringing residues A60 and A61 into close proximity with U52 and U53, while displacing U56 and U57 away from the duplex. In contrast, in the central region where the triple pairing occurs, no major differences are observed compared with the conventional MD simulations, apart from minor local shifts.

**Figure 3.**
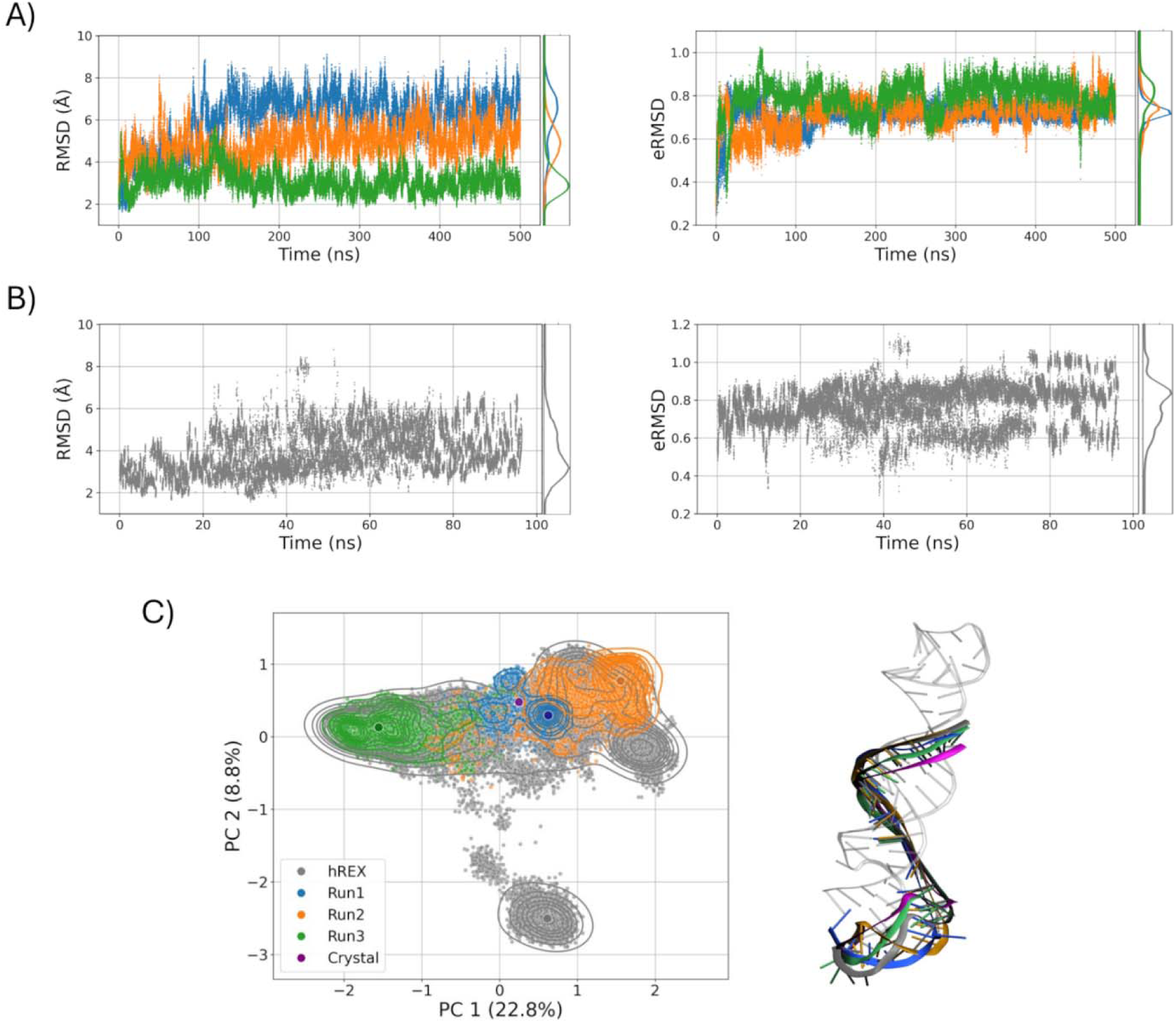
Conformational dynamics of MALAT1 in conventional and enhanced MD simulations. A) Timeseries and distribution of RMSD and eRMSD computed on the poly(A) residues using the crystallographic conformation as a reference in A) three independent conventional MD simulations and B) the HREX MD simulation. C) PCA of the aggregated conformations sampled in the conventional and HREX MD simulations, color-coded consistently with panels A-B) and with the crystallographic conformation indicated in purple. The circles indicate the representative conformations of the poly(A) tail from the most populated clusters of each simulation. On the right, the representative conformations are superimposed on the crystal one using the same color scheme.

### Pocket Communication Analysis Reveals Potential Binding Sites

Sampling a broad conformational space of the MALAT1 triple helix does not necessarily imply the presence of binding sites capable of accommodating the small molecules in the reference library, nor does it guarantee that the observed conformational variability is reflected in pocket diversity. To address these issues, we carried out a systematic pocket identification and pocket communication analysis using the Pocketron algorithm. The analysis yielded a total of 37 pockets, encompassing those present in the initial structure and those dynamically formed during the simulation. A comprehensive picture of pocket communication within the MALAT1 structure is reported by the 37×37 correlation matrix derived from the Pocketron analysis (**Fig. S3**), where each element quantifies the communication between corresponding pocket IDs (pIDs). These pockets can be visually represented as spheres (**Fig. 4A)**, with sphere radius proportional to pocket volume and color indicating pocket persistency (percentage of trajectory frames with non-zero volume).

**Figure 4.**
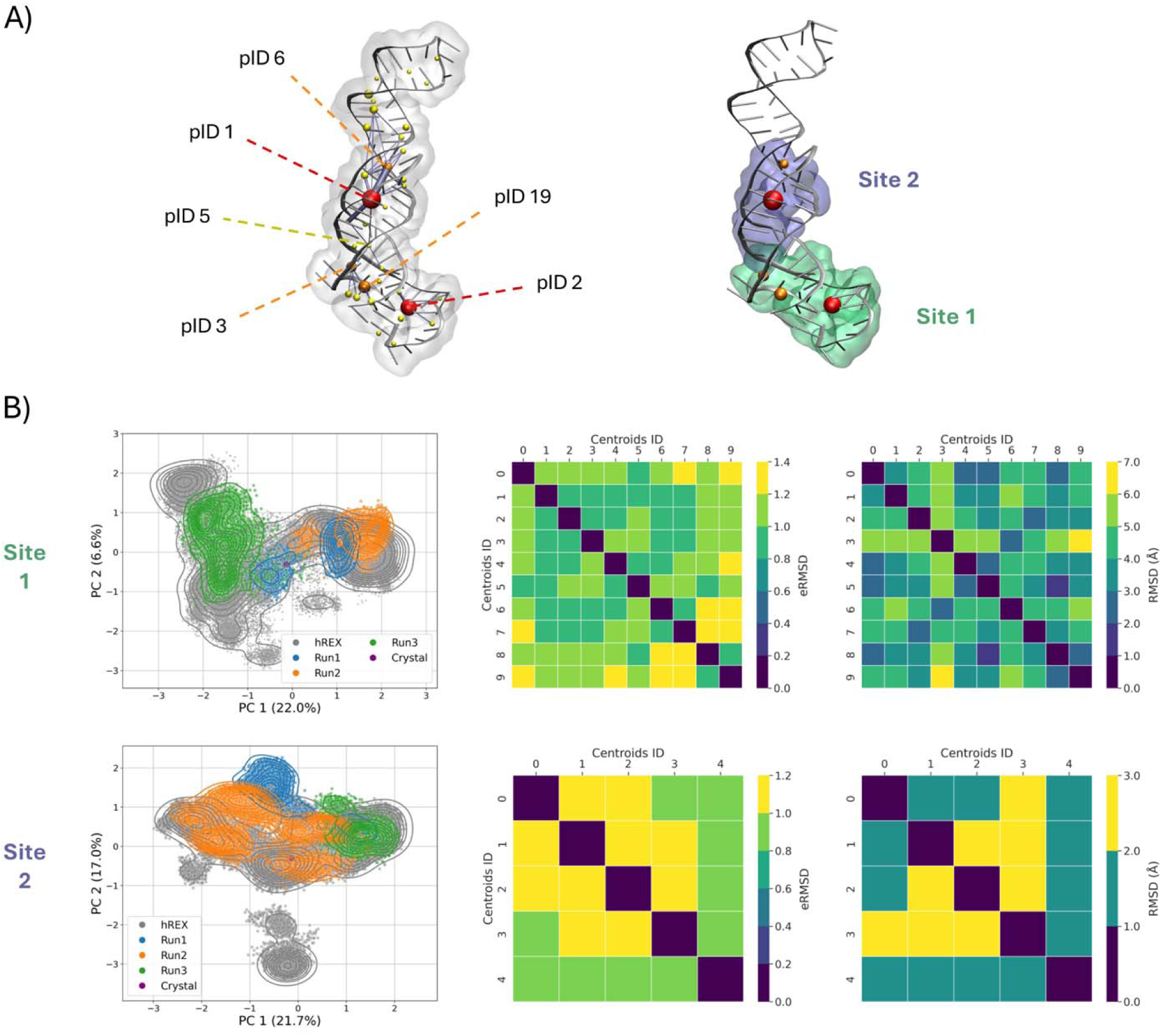
Generating a conformational ensemble of the promising binding regions. A) Pocket communication analysis on the standard MD simulations (left) identified two major interconnected regions in the MALAT1 structure, to use as promising binding sites for subsequent docking (right). Yellow, orange and red spheres correspond to pockets displaying increasing persistence, with sphere size reflecting the pocket volume, and edge thickness proportional to communication strength. B) PCA of the combined MD simulations (left), and eRMSD and RMSD matrices of most dissimilar cluster centroids from the HREX MD simulations (right), performed separately for the two binding regions.

The color scheme corresponds to the following persistence ranges: yellow (<33%), orange (33-66%), and red (>66%). Notably, the most persistent pockets also exhibit larger volumes, suggesting their potential suitability for further analysis and targeting in drug discovery efforts. To further assess the suitability of these pockets for ligand binding, we employed Pocketron to track residue exchanges between pockets and quantify the extent of communication across different target regions. This involved calculating the average number of “merge” and “split” events,(*74, 85*) which can be then visualized in a 3D graph (**Fig. 4A)**, with edge thickness proportional to the strength of communication between pockets. Overall, the analysis reveals two promising regions that harbor the largest and most persistent pockets. In the lower portion of the triple helix, a localized exchange of residues around the high-volume, high-persistence pocket 2 can be observed. In the adjacent region, another short communication pathway is present, including pockets 19 and 3, displaying a slightly lower persistence. Interestingly, a more long-range pathway originating from pocket 19 can also be observed, traversing pockets 3 and 5 to reach the second high-volume, high-persistence pocket 1, in the upper portion of the structure. Based on these insights, we next focused our analysis on the two regions comprising the most persistent pockets (**Fig.4A, Tab. 1**). These regions, hereafter named Site 1 (including pIDs 2, 3, and 19, in the lower portion of the triple helix) and Site 2 (including pIDs 1 and 6, in the upper portion), represent promising areas for further exploration as ligand binding sites, in good agreement with the work of François-Moutal et al.(*86*) Notably, not only do the pockets engage in frequent residue exchanges with neighboring ones, but they also exhibit communication across the two regions, albeit to a lesser extent. This is interesting, as inter-site communication hints at potential allosteric effects(*87*) that could propagate through the structure and ultimately influence the stability of the entire triple helix.

**Table 1.** Features of the major pockets identified from the MD simulations, namely pocket ID, average volume, residues comprised in the pocket, pocket persistency, and corresponding binding region.

| pID | Average volume<br>(Å<sup>3</sup>) | Residues | Persistency | Site |
| --- | --- | --- | --- | --- |
| 1 | 455 | 9, 10, 11, 39, 40,<br>68, 69, 70 | 96% | 1 |
| 6 | 99 | 6, 7, 64, 65 | 42% |  |
| 2 | 297 | 1, 2, 3, 4, 50, 51,<br>52, 54, 55, 56,<br>57, 58, 59, 60,<br>61, 62 | 69% |  |
| 3 | 179 | 45, 47, 48, 64, 65 | 62% | 2 |
| 19 | 294 | 47, 48, 49, 50 | 35% |  |

### Characterization of a Site-Specific Conformational Ensemble

With the binding site definition established, we next characterized the conformational ensemble spanned by Site 1 and Site 2 in terms of the spatial arrangements of their constituent residues. In analogy with the previous analyses, we computed site-specific eRMSD and performed PCA (see Methods) for both the conventional and HREX MD trajectories. **Figure S4** illustrates the eRMSD time series for both sites, with values ranging from approximately 0.6 to 1.3. HREX-MD simulations expanded the conformational space accessible to the two regions beyond that sampled by the three independent conventional MD runs. This effect is particularly evident in the PCA analysis (**Fig. 4B**, left panels). Comparing the two sites, Site 1 maintained a base pairing pattern closer to the reference compared to Site 2 (**Fig. S4**), whose dynamics were mostly driven by rearrangements of the triple helix. Specifically, Site 1 spanned values between 0.7 and 1.1, whereas Site 2 explored higher eRMSD values, except for Run 3, which remained closer to the reference.

To identify representative conformations of the two sites, we performed cluster analysis. To this end, we applied the Quality Threshold clustering algorithm (*79*) to the eRMSD matrix of the residues comprised in the two regions. The overall conformational diversity is captured by the pairwise eRMSD matrices (**Fig. 4B**, center and right panels), with eRMSD values spanning the 0.7-1.4 range for Site 1 and 0.7-1.2 for Site 2, highlighting a greater conformational variability, and thus greater flexibility, for Site 1. Then, to maximize the structural variability of the representative structures, we only retained cluster centroids with eRSMD above the cutoff of 0.7. This resulted in two independent ensembles comprising 10 and 5 structures for Sites 1 and 2, respectively. Taken together, these curated ensembles captured the full range of conformational dynamics observed in the simulations and provide a minimal, yet representative, set of structures to support efficient downstream docking studies.

### Complementary Binding Modes by Distinct Docking Programs

In a recent work, a library of 21 congeneric compounds was experimentally tested against the MALAT1 triple helix, providing a reference set of small molecules.(*51*) The compounds share a diminazene scaffold, decorated with different substituents in the ortho, meta and para positions (**Fig. 1B**). Thus, we used the generated conformational ensembles for the two binding regions in the RNA to dock the 21 compounds, in an ensemble docking spirit.(*40, 88*) We also included the conformations of Sites 1 and 2 of the crystal structure of the RNA, yielding 17 combined conformations. Using AutoDock GPU and rDock, we generated 250 poses per ligand with each program (500 poses per ligand overall). This process was repeated for all representative structures of the two sites, resulting in 8500 docked poses for each compound. We first assessed the extent of agreement between the poses generated by the two programs by performing dimensionality reduction across all the poses for all the RNA target conformations within each site.

**Figure 5** shows the results of PCA on the Cartesian coordinates for 5500 poses of P13 across 11 RNA conformations for Site 1 and 3000 poses across 6 RNA conformations for Site 2, obtained with the two software for compounds P13, which has the largest number of heavy atoms and represents the most challenging case for accommodation within the target due to its marked steric hindrance. While the poses occupy overlapping regions in the PC space, rDock was able to generate binding modes not accessed by AutoDock GPU, particularly in Site1. This result underscores the value of using multiple docking programs to capture a broader spectrum of potential binding poses on the RNA target. For the smaller Site 2, the poses from both programs occupy similar regions in the PC space, indicating lower diversity between the generated poses. Overall, for both sites, the first PC distinguishes elongated binding modes along the major groove and more compact ones.

**Figure 5.**
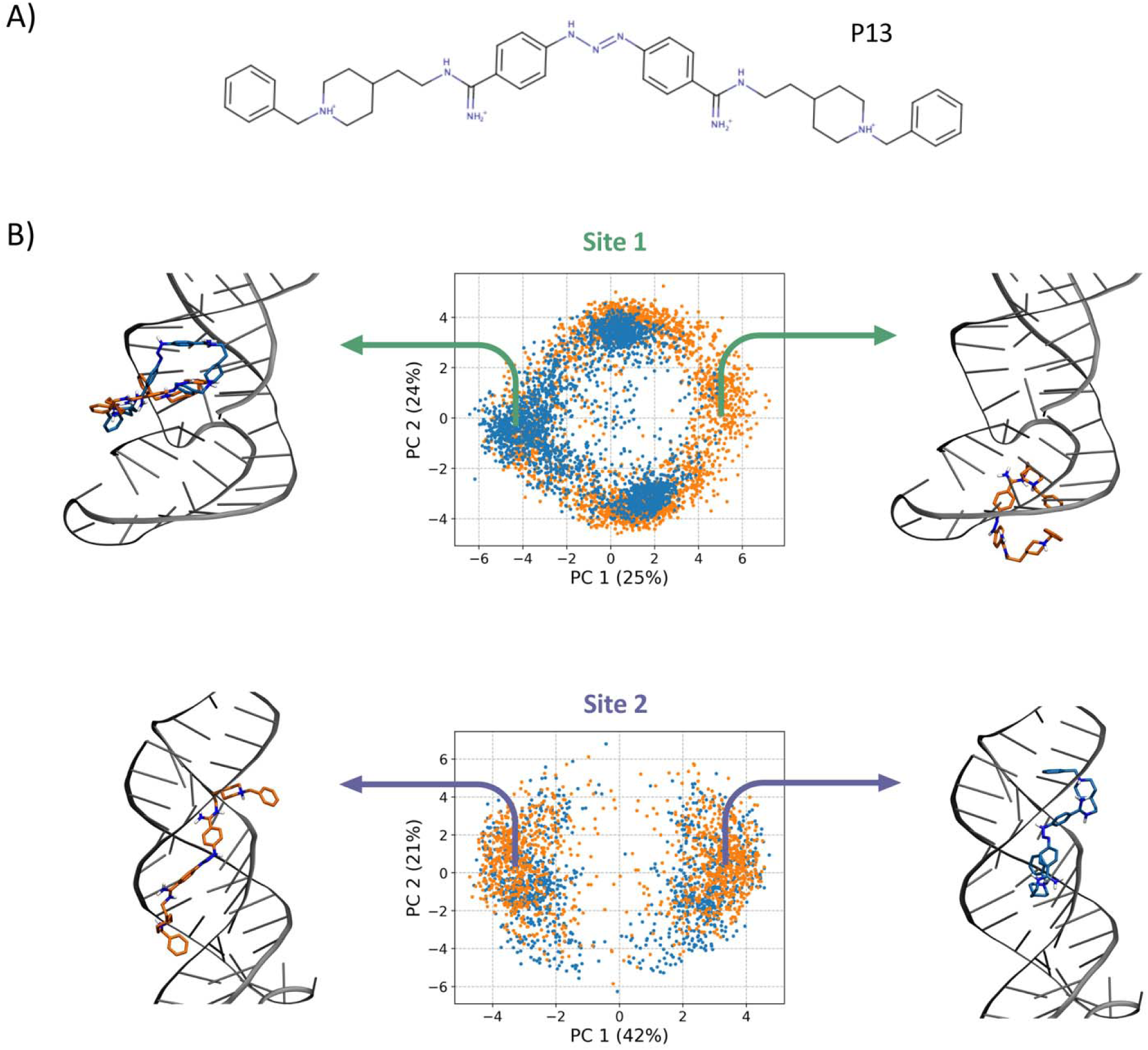
Binding poses generated for ligand P13. A) 2D structure of the diminazene analog P13. B) PCA on the aggregated poses generated for ligand P13 at the two sites, using the AutoDock GPU (blue) and rDock (orange) docking software. Representative binding poses from extreme regions along PC1 are shown, with ligand carbon atoms colored consistently with the software that generated the pose.

### Scoring Function Performance and Conformational Preference

To investigate how different scoring functions evaluate the same set of docked poses, we rescored all poses using multiple scoring functions, namely AutoDock, rDock, AutoDock Vina, AnnapuRNA, and SPRank. An overview of the results, covering all ligands and all combinations of docking programs and scoring functions for both Site 1 and Site 2, is reported in **Figures S5-S8**. The general trend that can be inferred is that poses tend to have more favorable scores when evaluated by the native scoring function of the program that generated them. For example, poses from AutoDock GPU scored overall better under the AutoDock scoring function, while poses from rDock performed better when evaluated by the native rDock scoring function (**Fig. S5 and S6**). This effect is more pronounced for rDock, which shows greater variability across the scores. AutoDock Vina exhibited a similar pattern to AutoDock, although to a lesser extent, suggesting it identifies stable poses across both docking programs more uniformly. By contrast, AnnapuRNA showed lower consistency (**Fig. S7** and **S8**), with poses generated by both programs largely overlapping within the same score range. However, several poses produced by AutoDock GPU received highly unfavorable scores, likely due to steric clashes. Interestingly, this behavior was absent when scoring rDock-generated poses, suggesting higher compatibility between the RNA-specific docking program and this scoring function. Nevertheless, this trend could not be generalized to all considered RNA-specific scoring functions, since SPRank exhibited the opposite behavior, assigning more favorable scores to AutoDock GPU poses in both binding sites.

These patterns are further illustrated in **Figure S9**, which shows the top-scoring poses for each ligand across all docking program-scoring function combinations. AutoDock and rDock scoring functions consistently favored their respective docking outputs for both sites. AnnapuRNA favored rDock-generated poses, while Vina showed a site-dependent behavior. In particular, for Site 1, Vina generally preferred rDock poses, though a few AutoDock GPU poses were also highly ranked. Conversely, for Site 2, Vina leaned towards AutoDock GPU poses. Similarly, SPRank favored AutoDock GPU poses for Site 2, while it identified top-scoring poses from both AutoDock GPU and rDock for Site 1.

To assess the effect of target conformation on docking outcomes, we compared the top-ranked pose from each conformational ensemble for every ligand (**Fig. S10**). Site 1 exhibited a broad distribution of best-scoring structures, with conformers 5, 7, 10, and the crystal structure frequently yielding the top scores across several scoring functions. AutoDock produced mixed results, favoring conformers 7, 10, and the crystal structure, whereas AutoDock Vina showed an even stronger preference for the crystal structure. rDock displayed a clear bias toward conformer 10, while ML-based scoring functions yielded more variable rankings: AnnapuRNA tended to favor conformers 5 and 10, and SPRank preferentially selected conformers 5 and 7. In contrast, Site 2 displayed more consistent behavior, with fewer conformers dominating (**Fig. S11**). When considering the best score per ligand irrespective of the target conformation, both AutoDock and Vina most frequently favored conformers 1, 3, and the crystal structure, with Vina showing a marked preference for the latter even at Site 2. Conformer 4 consistently emerged as the preferred structure for rDock across all ligands, as well as for AnnapuRNA, though the latter occasionally favored conformer 1. SPRank generally preferred conformers 4 and 3, with minor deviations across individual ligands.

### Evaluation of Docking Scores Against Experimental Affinity Trends

Our goal for the ensemble docking exercise was to evaluate whether any combination of docking software and scoring function could reproduce the experimentally observed affinity trends among the ligands. In this respect, the overall picture is summarized in **Figure S11**, which reveals that no clear correlation with experimental binding affinities could be established, except for AutoDock to a limited extent. For the latter, higher-affinity ligands generally received more favorable scores than lower-affinity ones, suggesting that AutoDock scores may be suitable in distinguishing binders from non-binders.

Interestingly, different from other scoring functions, AutoDock results displayed highly similar scoring patterns across the ligand set for both binding sites. This implies that the AutoDock scoring is largely independent of the specific binding site, highlighting a potential limitation in its ability to capture site-specific interactions for this case. The comparable scoring behavior across sites may stem from the chemical composition of the two binding regions, both featuring A-U base pairs with two G-C pairs in the middle. This similarity in composition can be particularly relevant for force-field-based scoring functions such as AutoDock, which heavily rely on atomic partial charges and van der Waals interactions.

Although AutoDock showed the most pronounced site-independent behavior, similar but less marked scoring patterns emerged for Vina and SPRank. Particularly interesting is the similarity observed within Site 1 for AnnapuRNA and SPRank. This comparable trend indicates that certain physicochemical properties of the ligands themselves may drive scoring outcomes across different methods. Overall, compared to AutoDock, AnnapuRNA and SPRank preserve some degree of site-specificity in their evaluations.

### Interaction-Based Interpretation of Ligand Affinity

To rationalize the observed variation in experimental binding affinities based on the predicted binding modes, we focused on ligand poses generated by AutoDock GPU, as its scoring function was the only one to exhibit a discernible correlation with experimental trends, albeit with the limitations discussed above. In particular, we analyzed ligand interactions within Site 1 and Site 2 using the recently developed fingeRNAt.(*40, 80*) The tool inspects complexes between nucleic acids and ligands to detect and classify non-covalent interactions. All ligand poses across the conformational ensemble were evaluated to generate an average interaction profile. As shown in **Figure S12**, the average interaction profiles were broadly similar between the two sites due to their comparable chemical features. A notable exception involved lipophilic interactions: Site 2 showed a sharp peak at approximately 5 Å, while Site 1 exhibited a bimodal distribution with peaks around 4.5 Å and 5.5 Å. To detect dominant interaction patterns among ligand poses, we further applied PCA. Although differences between sites were subtle, a clear trend emerged along PC1. Specifically, poses derived from low-affinity ligands were predominantly located in the negative region of PC1, whereas those from high-affinity ligands clustered toward positive PC1 values, indicating a separation in interaction patterns consistent with binding strength. In particular, analysis of the PC loadings revealed that positive values along PC1 are mainly associated with hydrogen-bond (distance range 2.25–3.88 Å) and cation–anion (2.23–5.45 Å) interactions. By contrast, lipophilic contacts (2.75–3.95 Å) are primarily associated with poses located on the negative side of PC1. All other interactions and distance ranges display no significant contribution in the loading profile (**Fig. S13**).

To improve the interpretability of results, we performed a cluster analysis on the ligand binding poses. To discriminate major configurations of the ligands, we focused on the common diminazene scaffold of the 21 compounds. For this stage, we only retained the most populated clusters, covering about 50% of poses for further analysis (**Fig. 6A**). PCA of these selected clusters (**Fig. 6B**) showed improved separation of ligands by affinity, and the variance explained by the first two principal components increased from about 14% to approximately 26%, underscoring the effectiveness of this refinement.

**Figure 6.**
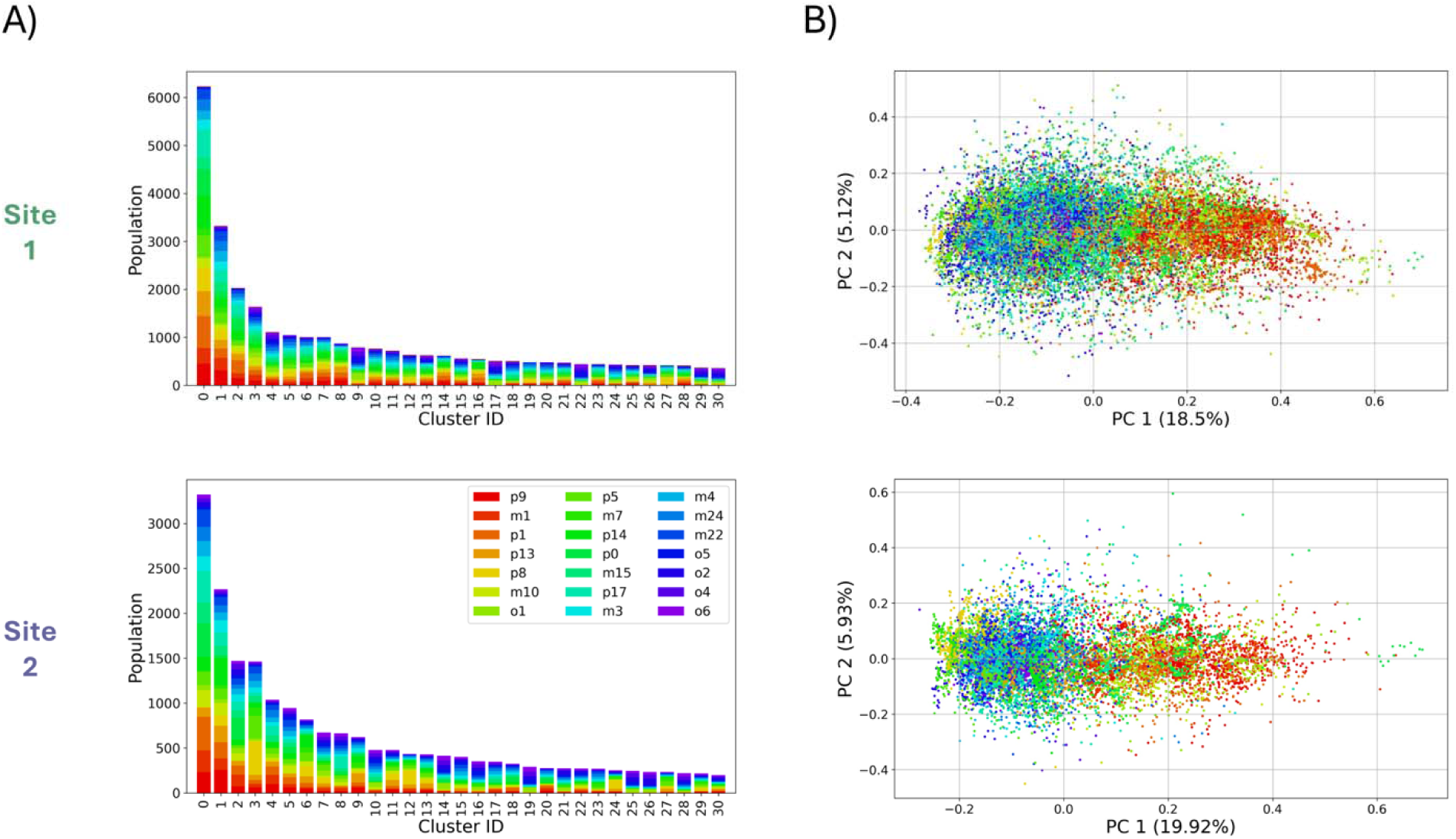
Characterization of the aggregated binding poses generated for all ligands. A) Cluster analysis and B) PCA were performed separately on Sites 1 and 2 (top and bottom panels, respectively), using about 50% of the total number of poses. Cluster populations are highlighted by occurrence per ligand, and the same color code is used for the PCA projections, ranging from red (highest relative affinity) to violet (lowest relative affinity).

To further explore affinity-related binding patterns, we analyzed the distribution of PC values within each cluster. For each ligand, we calculated the average PC1 score per cluster (**Fig. S14**–**S17**). Cluster 3 of Site 2 showed the strongest association with experimental binding affinities (R² = 0.6), suggesting that poses in this cluster, which feature a distinctive and consistent binding mode of the central scaffold, are most effective at distinguishing strong from weak binders.

A closer inspection of the binding poses within this cluster, focusing on ligands P9 (highest experimental affinity), P0 (unsubstituted scaffold), and O5 (lowest affinity), provides additional insight into the observed structure–activity relationships (**Fig. 7**). Starting with P0, which represents the common scaffold, the central benzene rings establish lipophilic contacts with the RNA bases, while the diminazene group forms hydrogen bonds with nearby residues. The amidine moiety at the opposite end further participates in hydrogen bonding with bases and cation–anion interactions with the phosphate backbone via its positively charged nitrogen atoms. The binding mode of P9 closely resembles that of P0 in terms of scaffold and amidine interactions. However, its nitrogen-rich, positively charged substituents enable additional hydrogen bonds and cation-anion contacts with the RNA phosphate groups. This expanded polar interaction network corresponds to the shift of P9 poses toward higher PC1 values in the PCA plot. By contrast, O5, which bears highly lipophilic substituents, predominantly forms lipophilic contacts with the aromatic bases, resulting in lower PC1 values. Overall, the experimental affinity trend can be rationalized based on the substituents’ charge and hydrogen-bonding capacity: nitrogen-rich, positively charged groups generally enhance affinity relative to P0, whereas purely aromatic or hydrophobic substituents reduce it, with the only exceptions being P5 and M7.

**Figure 7.**
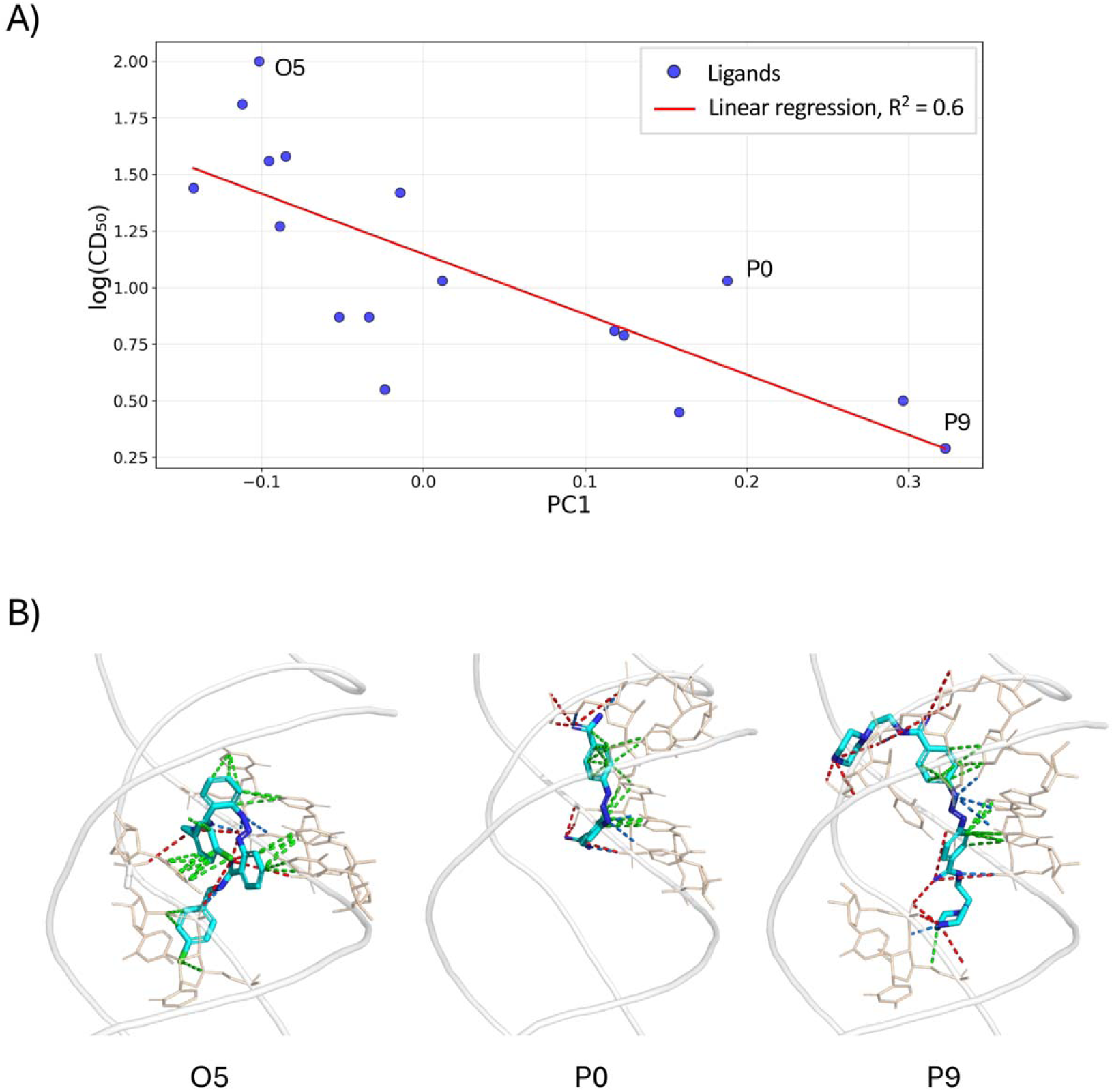
Rationalizing ligand affinity by predicted binding mode. A) Linear relationship between relative affinity and average PC1 score for the entire set of compounds. B) Proposed binding modes for representative ligands, namely the highest-affinity one (P9), the unsubstituted scaffold (P0), and the lowest-affinity ligand (O5). The RNA backbone is shown in tubes, while RNA nucleotides directly interacting with the ligands are displayed in licorice. Colored dashed lines indicate RNA-ligand interactions, with green, red, and blue representing lipophilic, cation-anion, and hydrogen bond interactions, respectively.

## Conclusion

In this study, we applied a comprehensive structure-based drug discovery (SBDD) pipeline to investigate how a congeneric series of diminazene derivatives interacts with the MALAT1 triple helix, a prototypical structured lncRNA with known druggability. Our approach integrated enhanced molecular dynamics simulations, pocket communication analysis, ensemble docking, and rescoring with both classical and RNA-specific methods. The simulations revealed two persistent and ligand-accessible binding regions (Site 1 and Site 2), while the pocket communication analysis highlighted a degree of dynamic coupling between RNA subregions, hinting at potential for allosteric modulation. Ensemble docking across the generated RNA conformational ensemble expanded the diversity of accessible binding poses, particularly within Site 1, and uncovered complementary binding subpockets depending on the docking tool used. Despite limitations, AutoDock achieved modest discrimination between high- and low-affinity ligands. Crucially, our analysis of clustered interaction patterns using fingeRNAt and principal component analysis offered a clearer view of structure–affinity relationships. One specific subpopulation of poses (Site 2, Cluster 3) showed clear trends in relation to experimental affinity, suggesting this as a potentially relevant binding mode. Nevertheless, several limitations emerged that mirror challenges consistently reported in the field. The accuracy of scoring functions, both general-purpose and RNA-specific, including machine learning–based approaches, proved insufficient for reliably predicting binding affinities across structurally similar ligands. This underscores the known sensitivity of scoring functions to initial target conformations and the inherent difficulty in accounting for ligand and target flexibility, especially in nucleic acid systems. Moreover, our results reinforce prior observations that accurately generating and evaluating docking poses for RNA targets remains an open challenge, further complicated by the lack of high-resolution experimental binding affinity data for nucleic acid-ligand complexes. These data gaps hinder the development, training, and validation of more robust predictive models. Additionally, although the RNA model used here was structurally resolved and subjected to extensive conformational sampling, it may not fully capture the plasticity and environmental complexity present in cellular contexts. The absence of experimental confirmation for the predicted binding poses, combined with the limited chemical diversity of the ligand series, also limits the generalizability of our findings.

Altogether, our work underscores both the potential and the limitations of current SBDD strategies when applied to dynamic RNA targets. While advanced computational pipelines can extract useful structure–activity relationships and identify plausible binding modes, achieving predictive reliability remains out of reach. Progress in RNA-targeted drug discovery will require not only better scoring functions and more flexible modeling strategies but also broader access to high-quality experimental data and integrative validation frameworks.

## ASSOCIATED CONTENT

### Supporting Information

Comparison of grid box and cavity mapping in AutoDock and rDock, respectively; inter-pocket communication matrix; eRMSD of the two sites in conventional and HREX MD; docking scores for all the generated poses on all RNA conformers of the two sites; analysis of RNA-ligand interactions via FingeRNAt; loadings of PC1 for different interaction features; PCA on the molecular fingerprints of all ligand poses for the two sites and resulting correlations.

## AUTHOR INFORMATION

### Author Contributions

The manuscript was written through contributions of all authors. All authors have given approval to the final version of the manuscript.

## ACKNOWLEDGMENT

We gratefully acknowledge the Data Science and Computation Facility and its Support Team at Fondazione Istituto Italiano di Tecnologia for computing time and support on the Franklin HPC system.

## ABBREVIATIONS

RNA: Ribonucleic acid
MALAT1: metastasis-associated lung adenocarcinoma transcript 1
MD: Molecular Dynamics
HREX: Hamiltonian Replica Exchange
SBDD: structure-based drug discovery.

