## Supplementary Material for "Critical Assessment of a Structure-Based Pipeline for Targeting the Long Non-Coding RNA MALAT1"

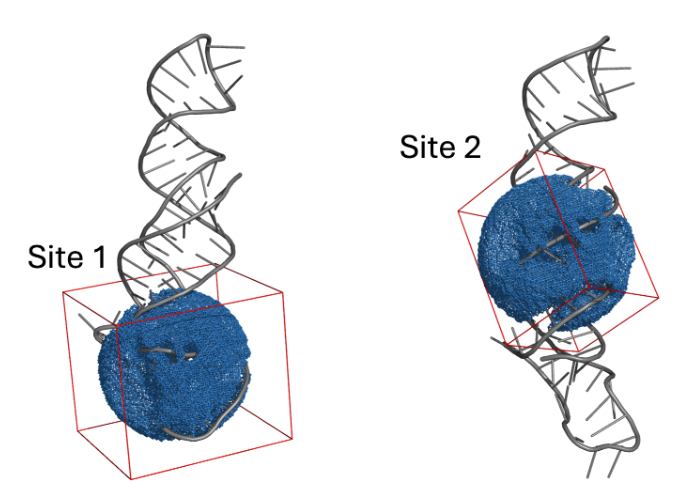

**Figure S1.** Comparison of AutoDock grid box (in red) and rDock cavity search (in blue) for the two binding regions.

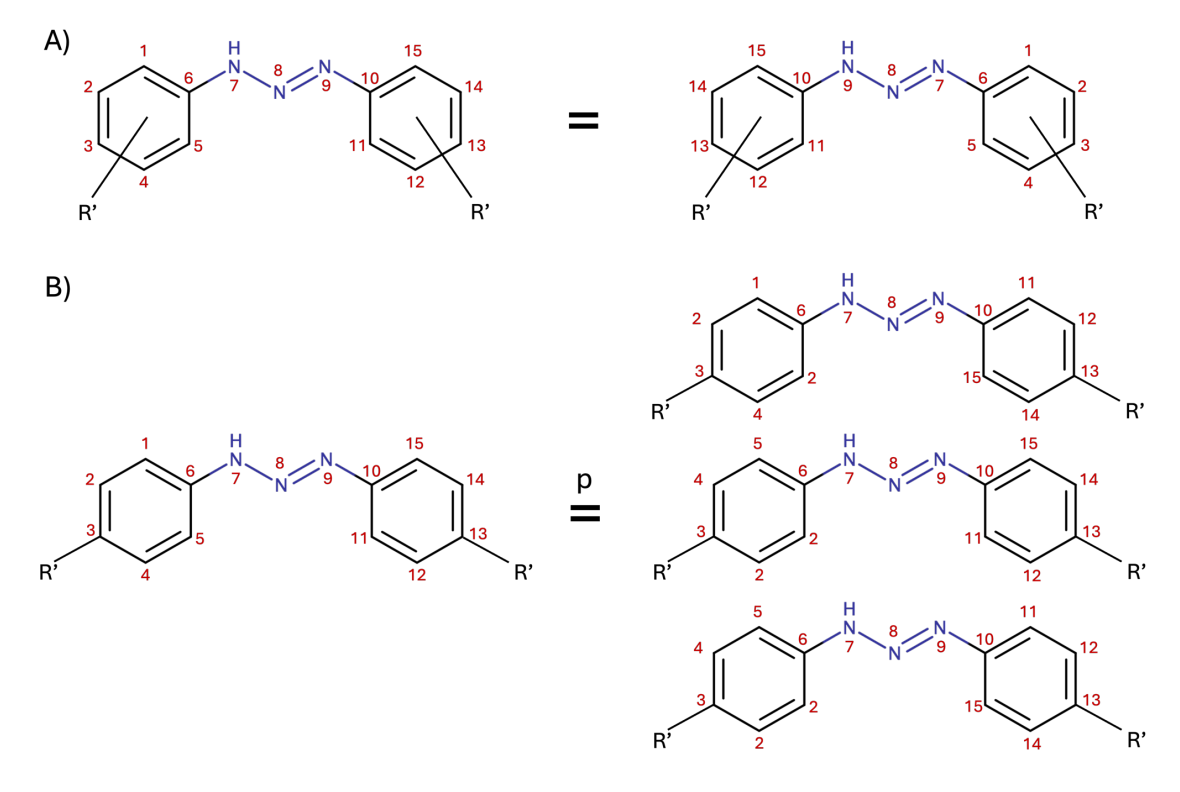

**Figure S2.** Atom index transformations used for minimum RMSD calculations. A) Symmetry operation applied to all ligands based on reflection across the σz plane passing through atom N8. B) Additional transformations specific to para-substituted ligands, leveraging their higher symmetry: rotations around the N9–C10 bond, the N7–C6 bond, and both simultaneously.

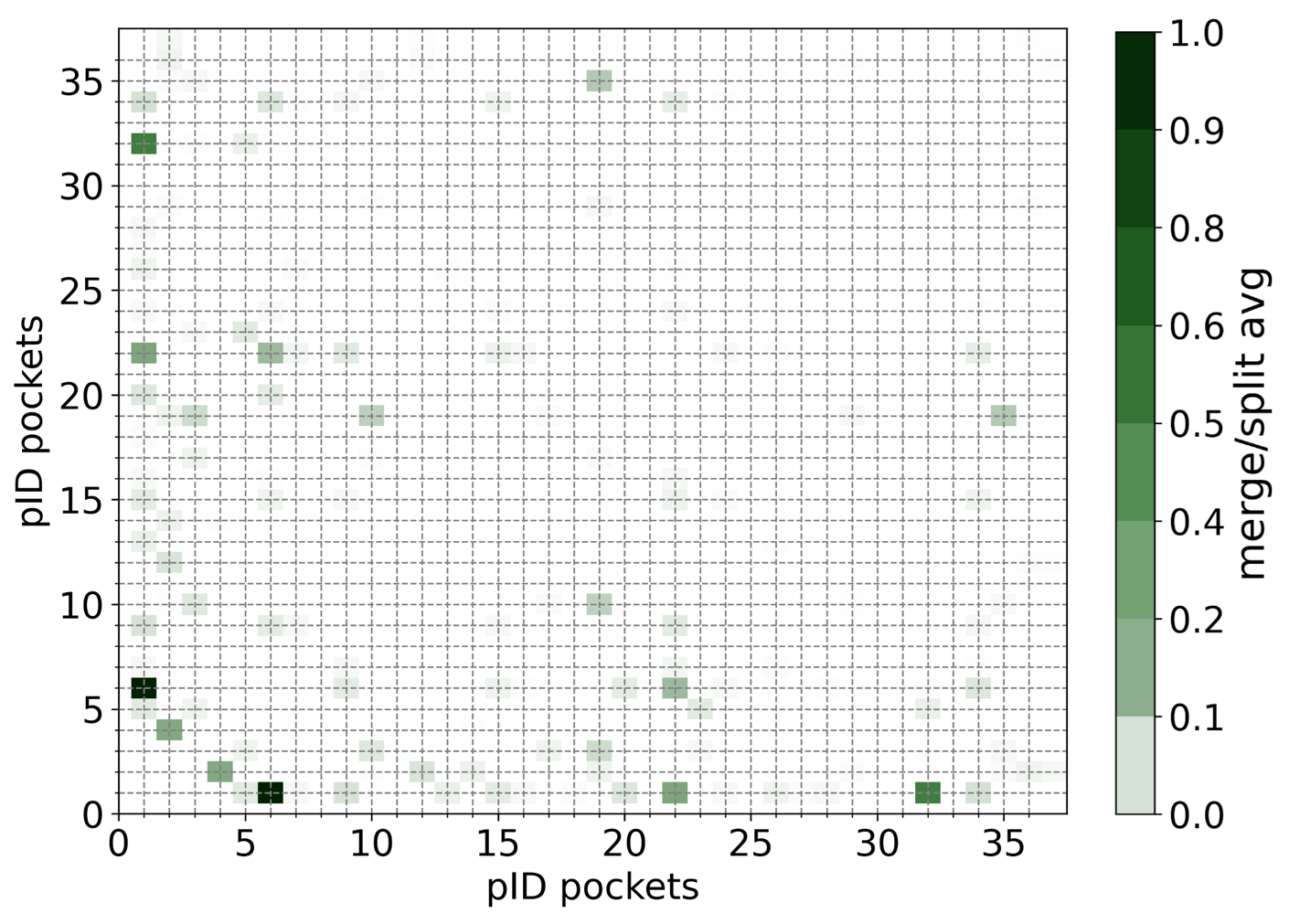

**Figure S3.** Inter-pocket communication matrix obtained from the Pocketron pocket communication analysis.

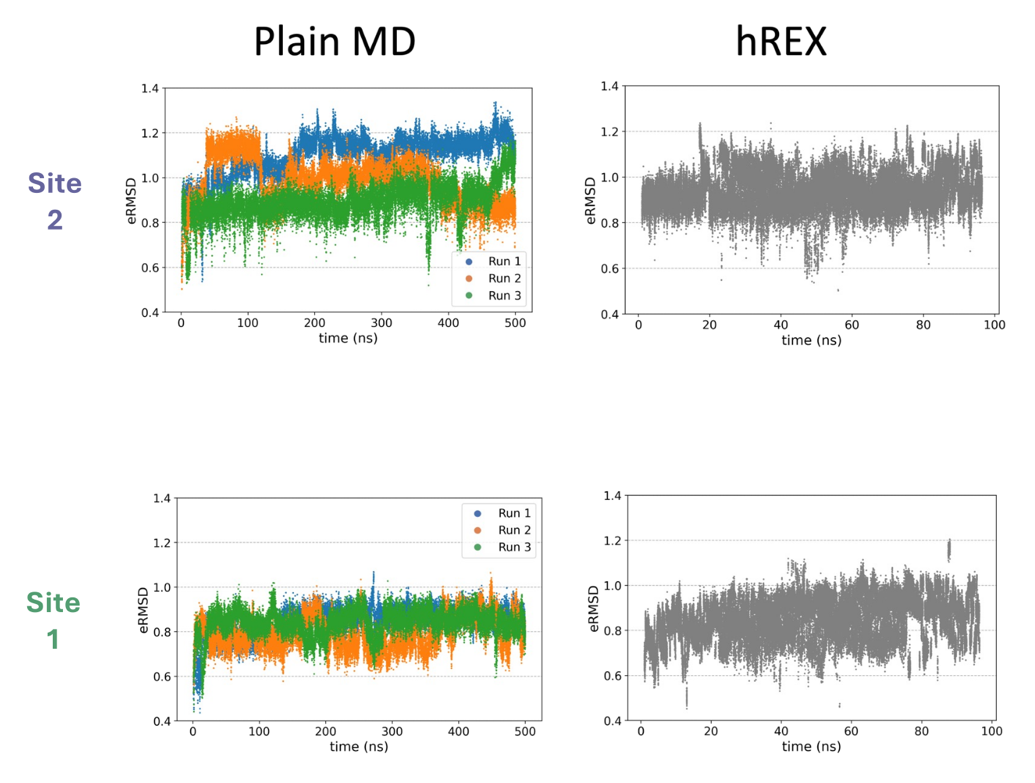

**Figure S4.** eRMSD of the plain (left) and hREX (right) MD simulations for the two binding regions.

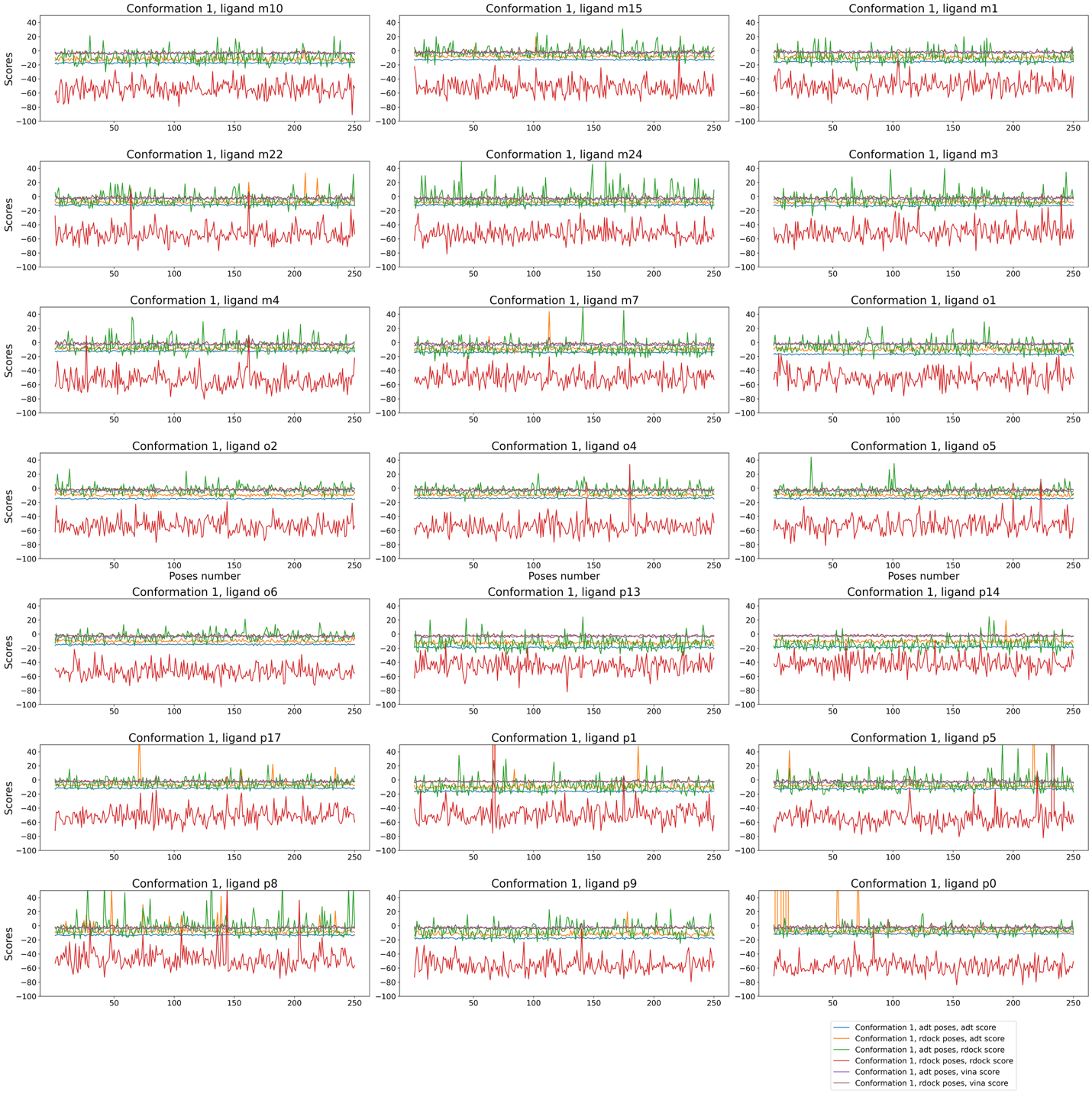

**Figure S5.** Docking scores for all the generated poses for a sample conformation of Site 1, color-coded by scoring function and docking software: Autodock (blue/orange), rDock (green/red), Vina (purple/gray). Blue/green/purple lines represent poses generated with Autodock GPU, while orange/red/gray lines represent those from rDock.

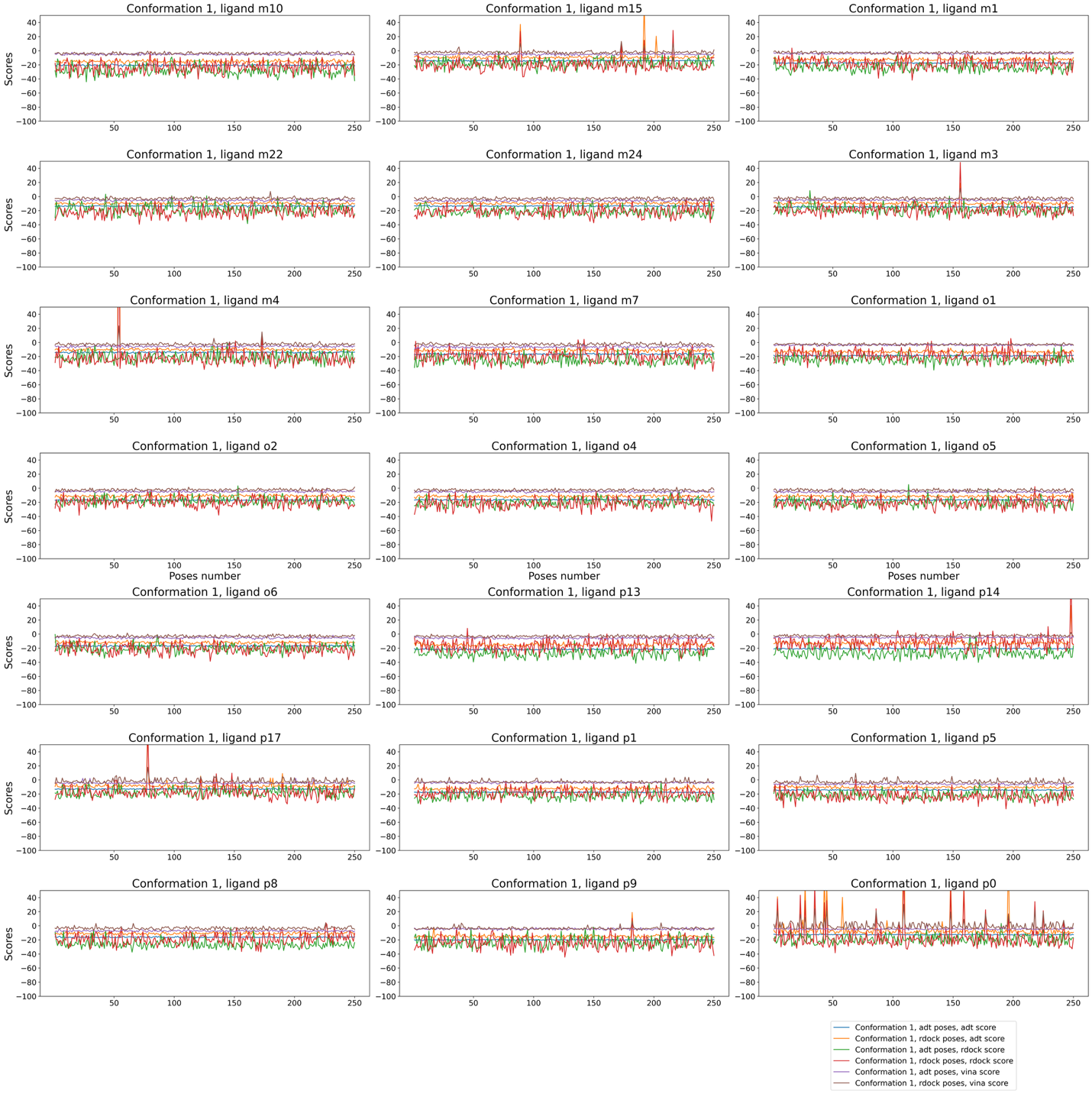

**Figure S6.** Docking scores for all the generated poses for a sample conformation of Site 2, color-coded by scoring function and docking software: Autodock (blue/orange), rDock (green/red), Vina (purple/gray). Blue/green/purple lines represent poses generated with Autodock GPU, while orange/red/gray lines represent those from rDock.

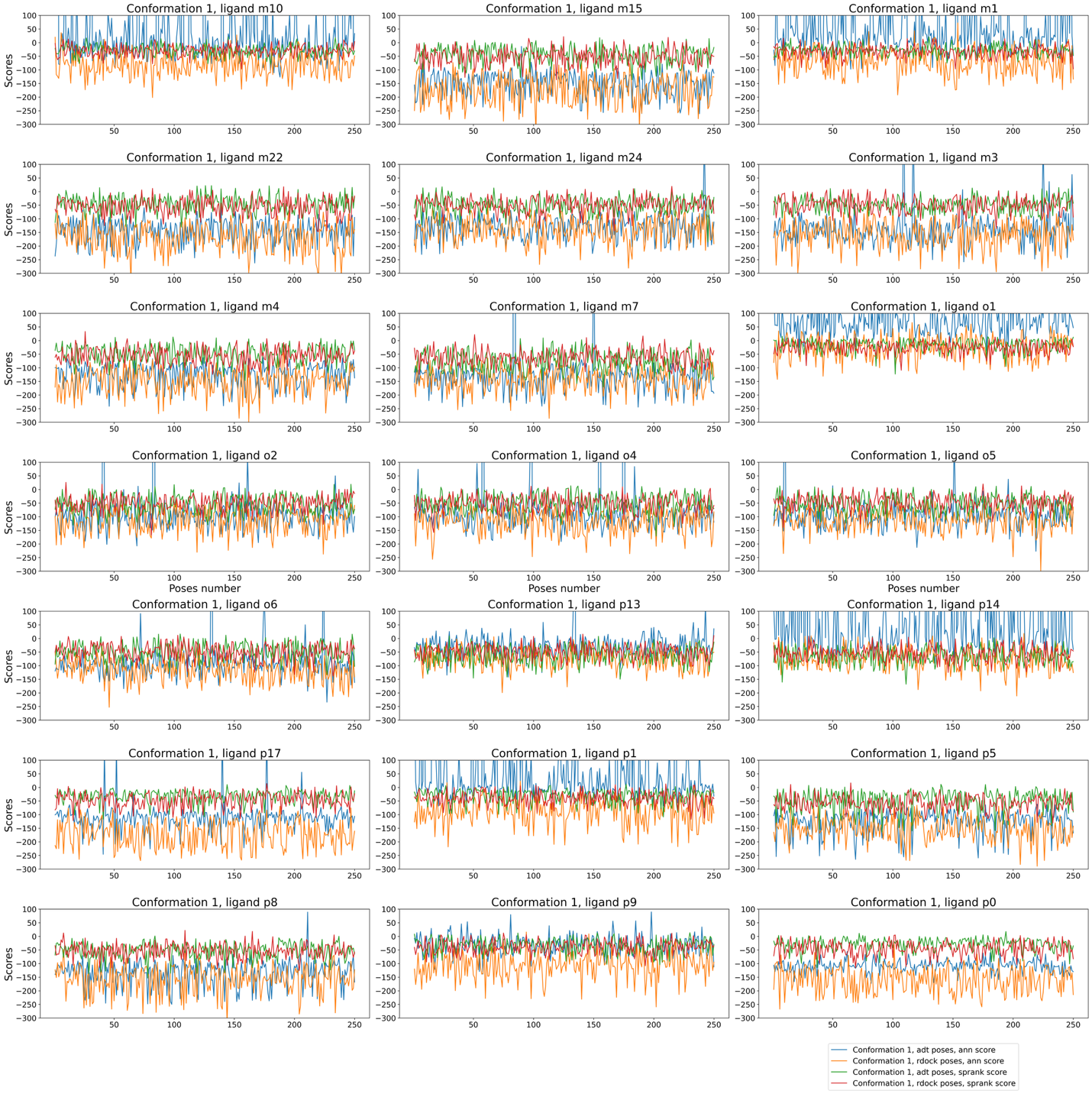

**Figure S7.** Docking scores for all the generated poses for a sample conformation of Site 1, color-coded by scoring function and docking software: AnnapuRNA (blue/orange) and SPRank (green/red). Blue and green lines represent poses generated with Autodock GPU, while orange and red lines represent those from rDock.

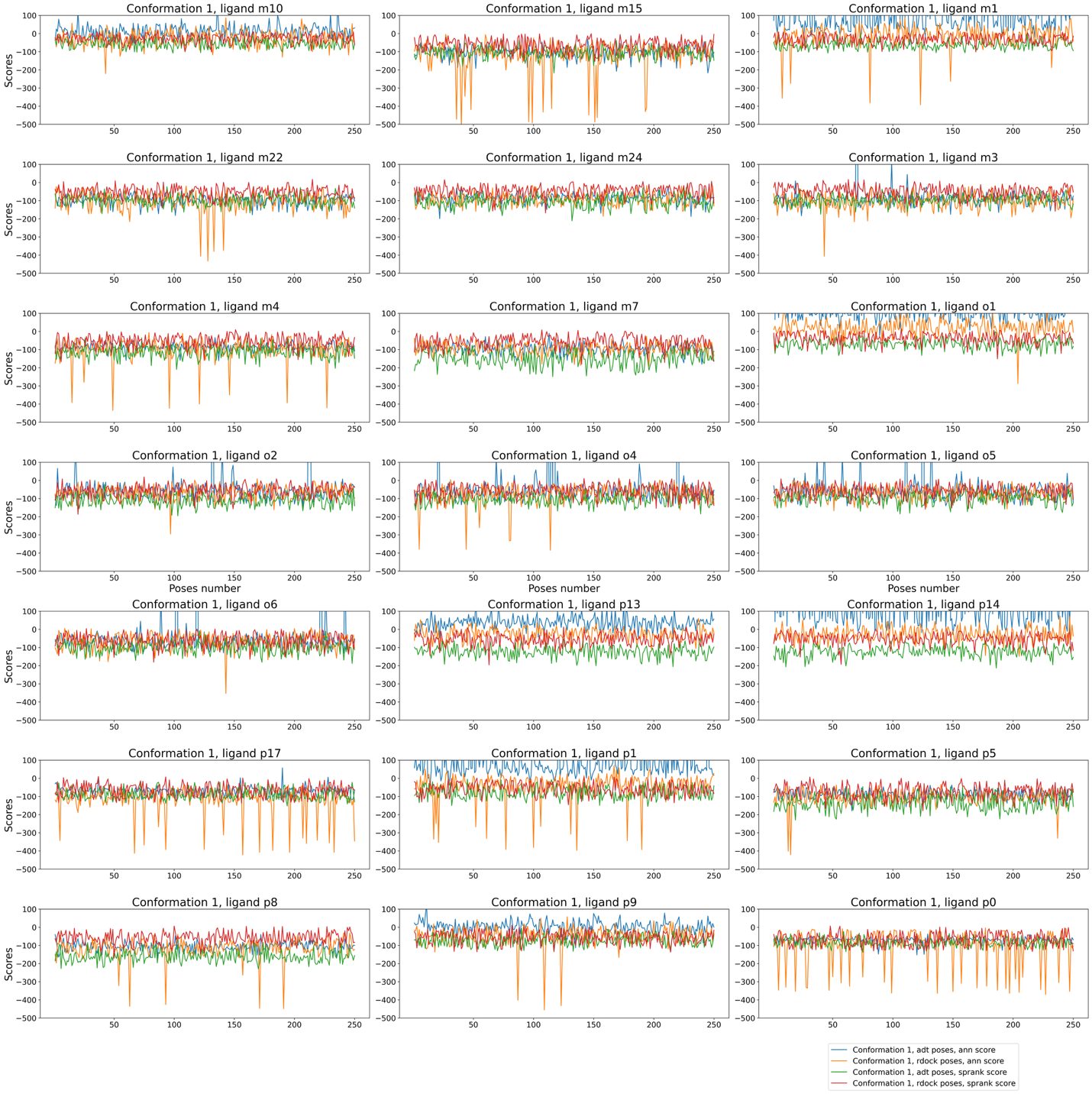

**Figure S8.** Docking scores for all the generated poses for a sample conformation of Site 2, color-coded by scoring function and docking software: AnnapuRNA (blue/orange) and SPRank (green/red). Blue and green lines represent poses generated with Autodock GPU, while orange and red lines represent those from rDock.

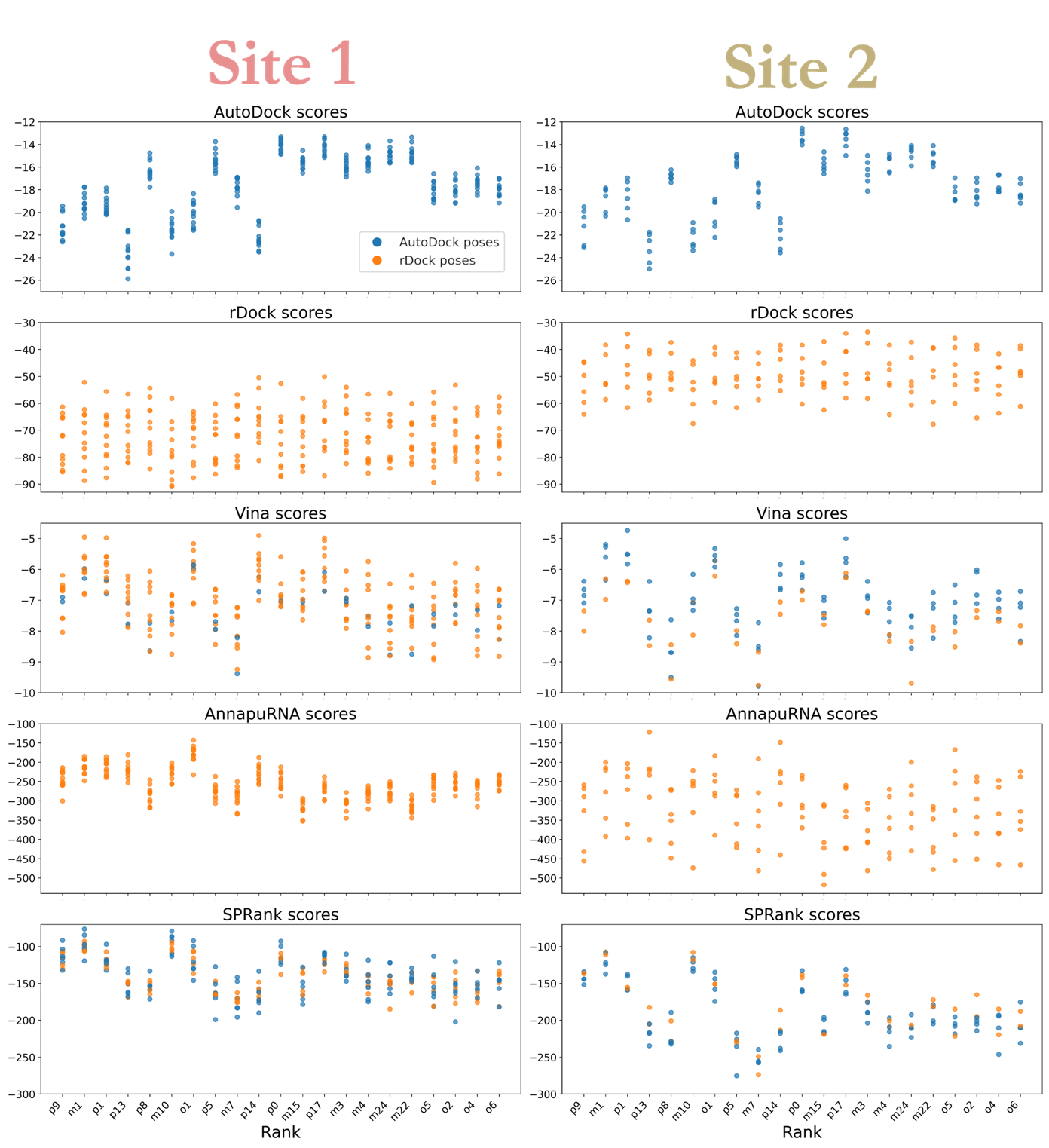

**Figure S9.** Evaluation of ligand-target binding using multiple scoring functions for both sites. Top-ranked poses for each conformer are color-coded by docking software: AutoDock GPU (blue), rDock (orange).

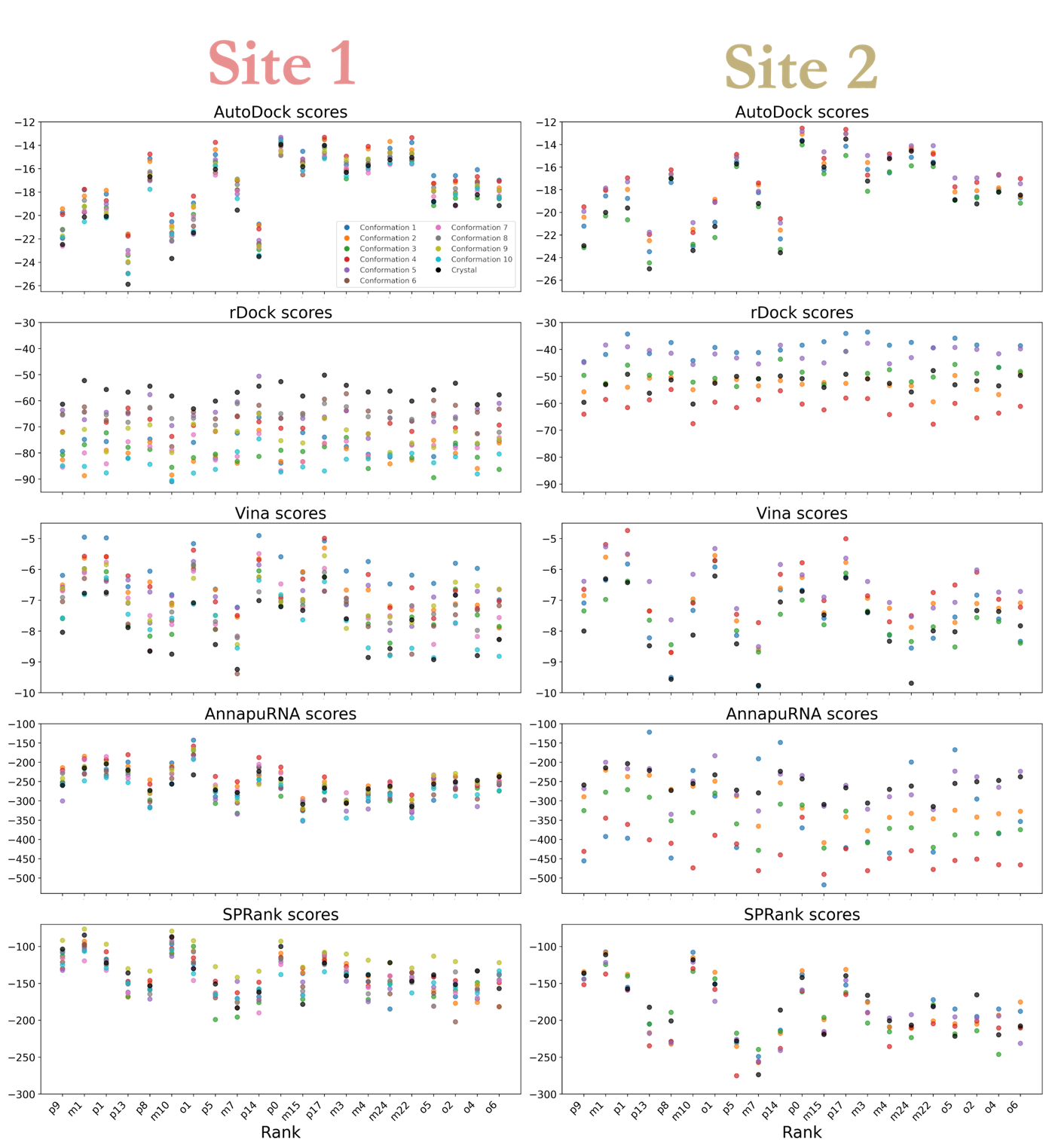

**Figure S10.** Evaluation of ligand-target binding using multiple scoring functions. Top-ranked poses for each target’s conformation with colors indicating their respective ensemble element for both sites.

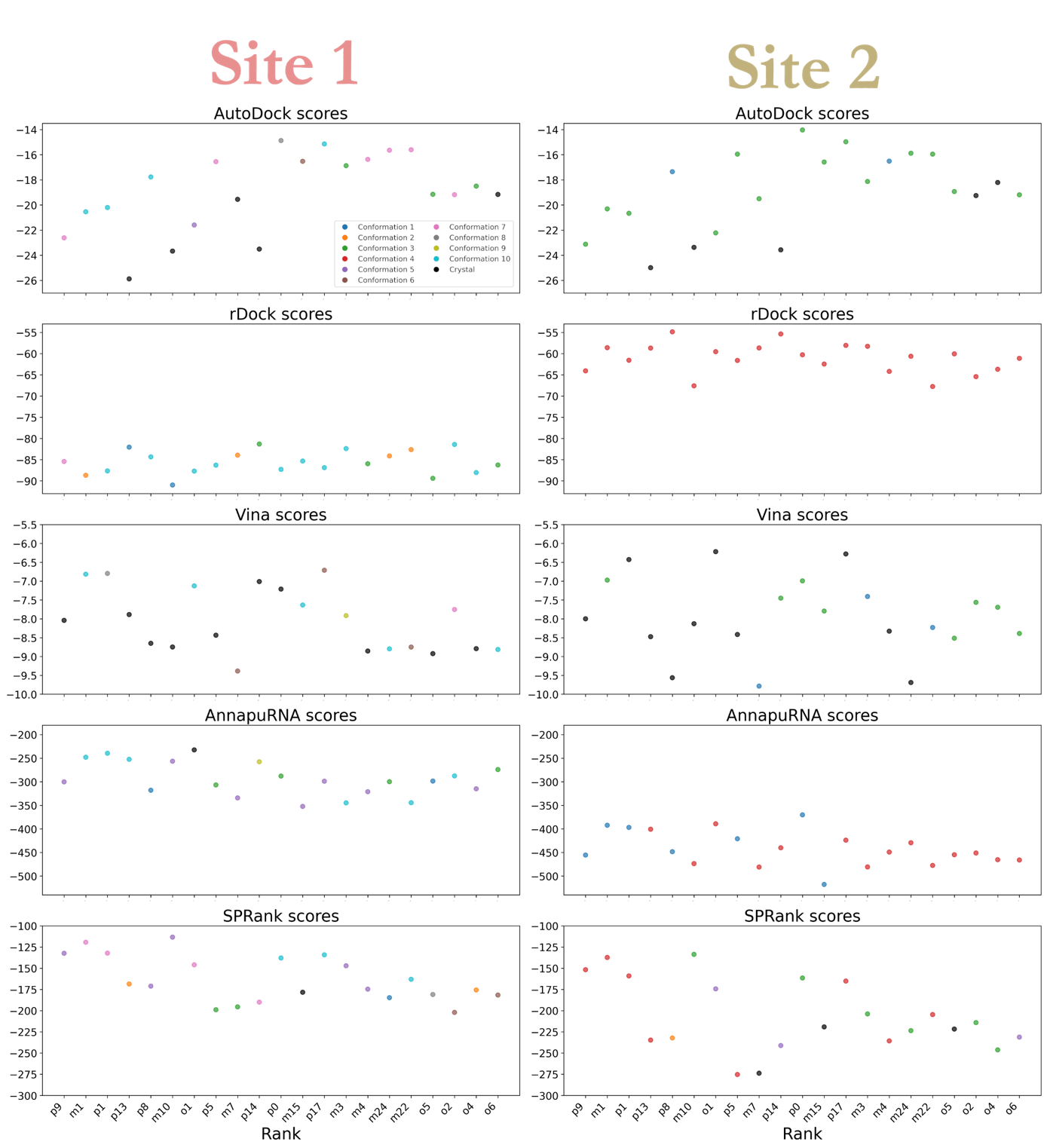

**Figure S11.** Evaluation of ligand-target binding using multiple scoring functions. Top-ranked pose for each conformation is extracted with colors indicating their respective element in the ensemble for both sites.

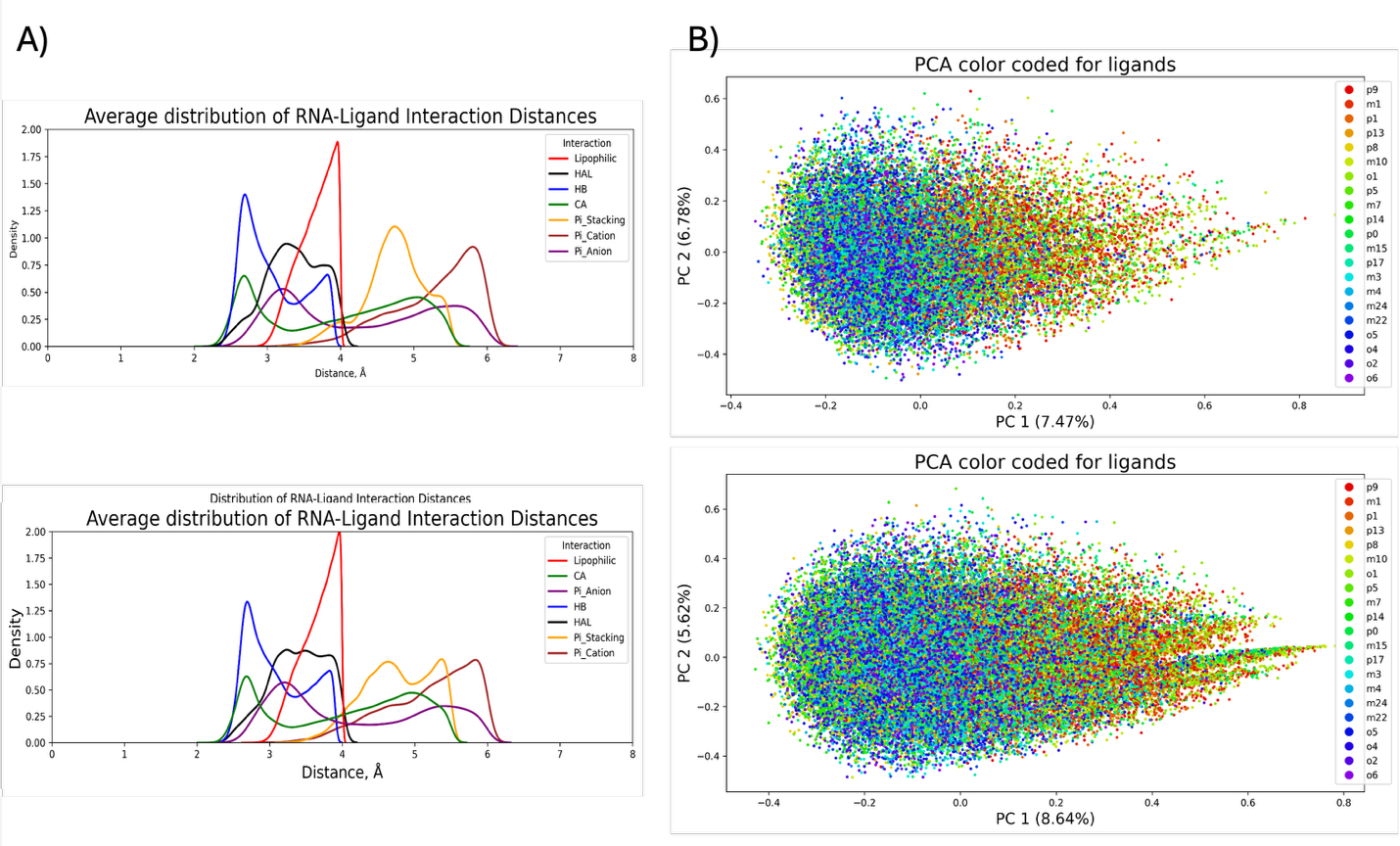

**Figure S12.** FingeRNAt analysis. (A) Average density distribution of intermolecular interactions between ligands and RNA. (B) Principal Component Analysis (PCA) depicting the distribution of all ligand poses for both binding sites.

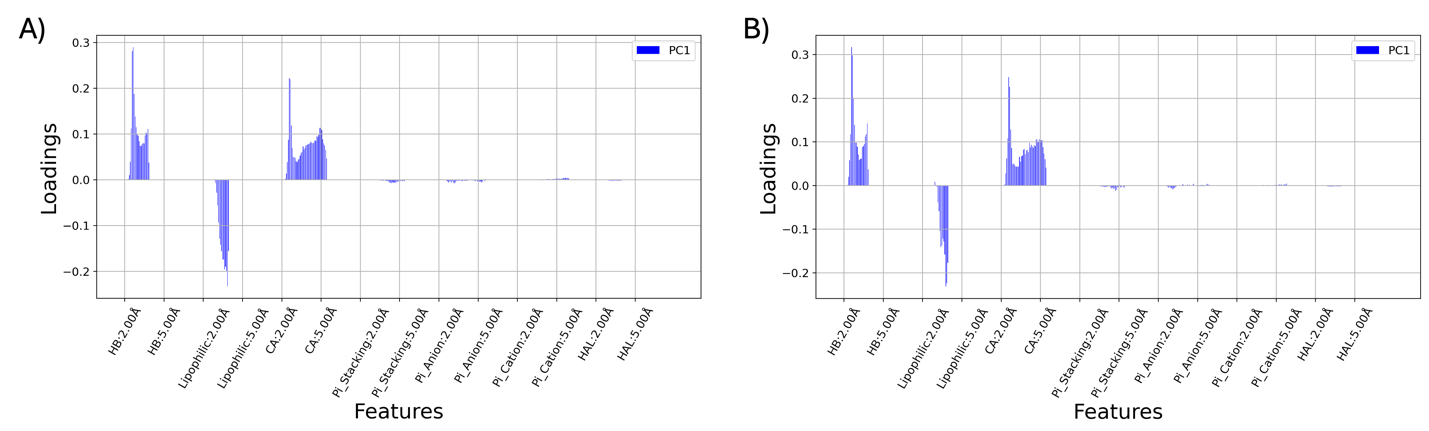

**Figure S13.** Loadings of the first principal component (PC1) for the interaction features in **(A)** Site 1 and **(B)** Site 2.

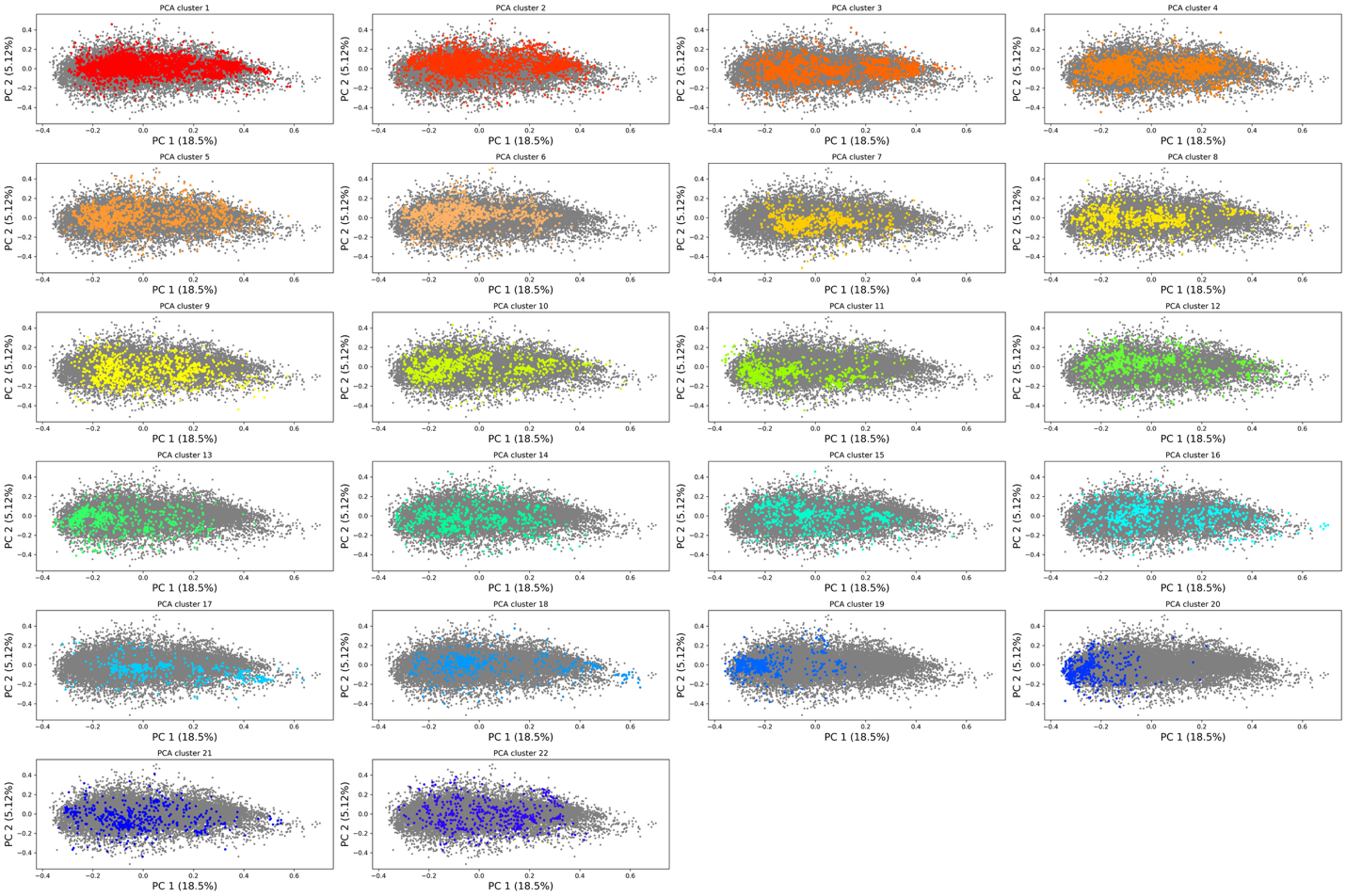

**Figure S14.** PCA performed on the molecular fingerprints of all ligand poses for Site 1. The gray dots represent the full dataset projected onto the first two principal components (PC1 and PC2). Each subplot highlights, in color, the data points corresponding to a specific cluster extracted from the clustering analysis.

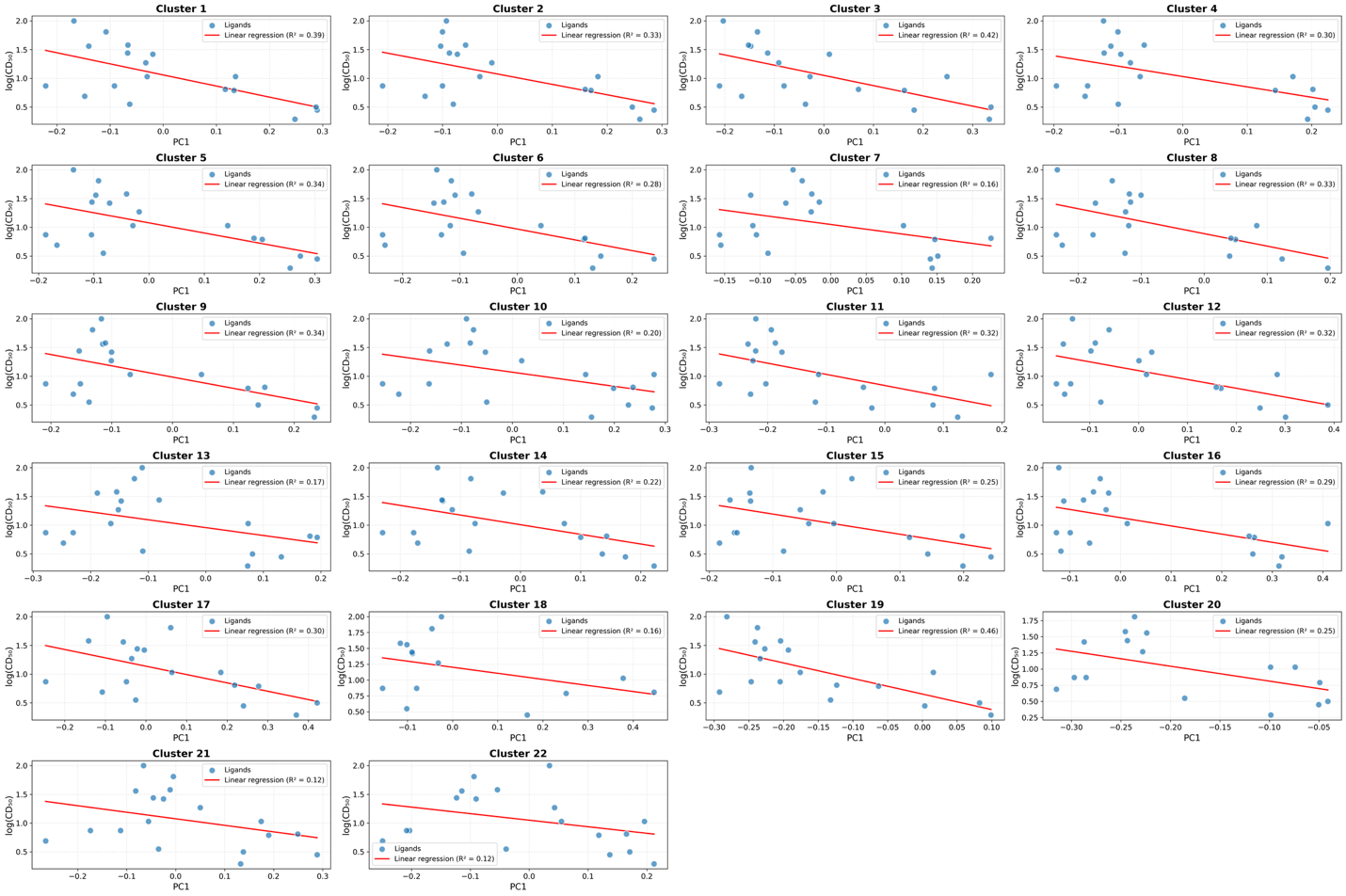

**Figure S15.** Linear relationship between the relative binding affinity and the average PC1 scores for each ligand across all identified clusters of Site 1. The red line represents the linear regression fit, with its equation and the coefficient of determination R^2^ reported in the legend.

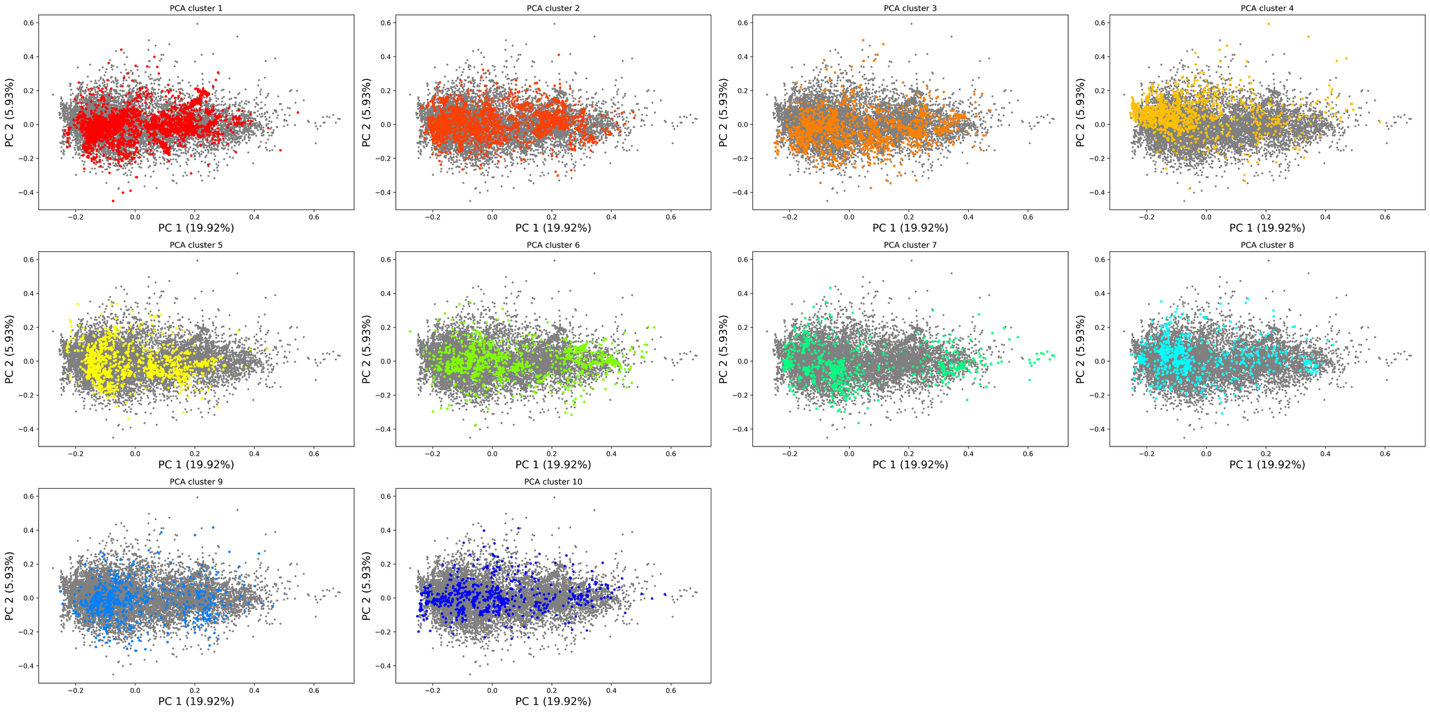

**Figure S16.** PCA performed on the molecular fingerprints of all ligand poses for Site 2. The gray dots represent the full dataset projected onto the first two principal components (PC1 and PC2). Each subplot highlights, in color, the data points corresponding to a specific cluster extracted from the clustering analysis.

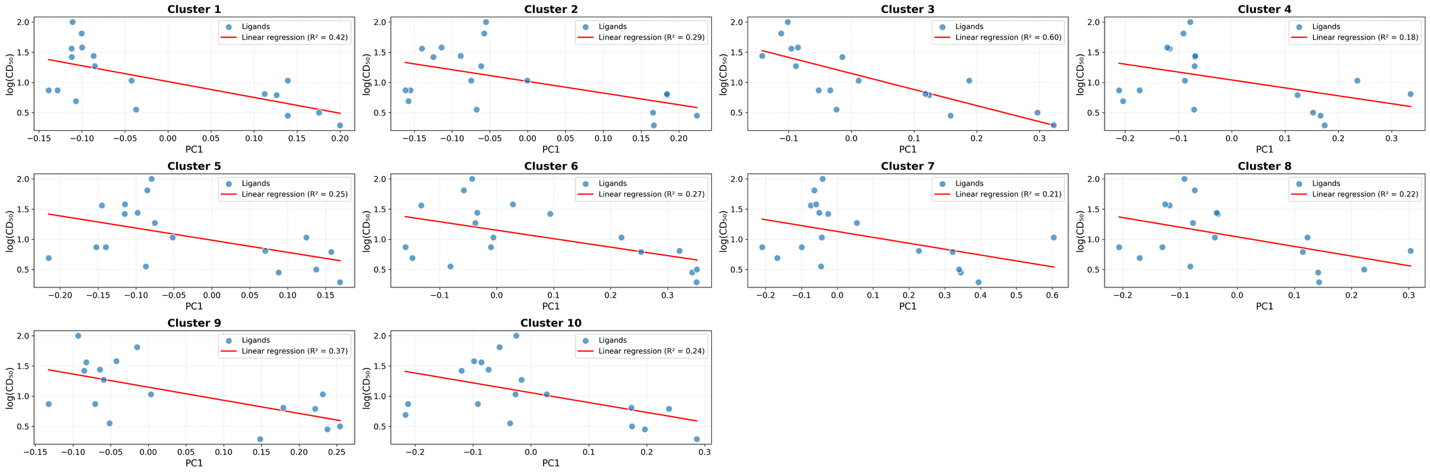

**Figure S17.** Linear relationship between the relative binding affinity and the average PC1 scores for each ligand across all identified clusters of Site 2. The red line represents the linear regression fit, with its equation and the coefficient of determination R^2^ reported in the legend.
